# ADAR1p150 RNA binding, independent of A-to-I RNA editing, buffers immunogenicity from the tonic type I IFN induced transcriptome *in vivo*

**DOI:** 10.1101/2025.10.14.682456

**Authors:** Jacki E Heraud-Farlow, Scott R Taylor, Ankita Goradia, Alistair M Chalk, Paul Harrison, Zhen Liang, Monique F Smeets, David J Izon, Shi-Bin Hu, Jin Billy Li, Louise E Purton, Carl R Walkley

## Abstract

ADAR1 edits adenosine to inosine (A-to-I) in double-stranded RNA to prevent MDA5 sensing of cellular transcripts, its key physiological role. However, editing-independent functions of ADAR1 remain poorly understood. Using a series of *Adar1* mutant mice rescued by loss of MDA5 and PKR, we investigated isoform-specific, editing-independent roles of ADAR1 *in vivo*. We found that the cytoplasmic ADAR1p150 isoform is essential for maintaining peripheral T cell numbers and differentiation of hematopoietic stem and progenitor cells (HSPCs). In bone marrow transplants, ADAR1p150 protein, but not its editing activity, was crucial for T cell regeneration and HSPC repopulation, demonstrating a cell-intrinsic function in hematopoiesis. Experiments with IFNβ-treatment of purified HSPCs *in vitro* and *in vivo* IFNAR1 neutralization revealed hypersensitivity to tonic type I interferon (IFN) in the absence of ADAR1p150. Using cell lines, we demonstrate that type I IFN activates the OAS-RNaseL pathway, leading to cell death. This study shows that tonic type I IFN induces immunogenic cellular RNAs in sterile conditions. ADAR1p150 suppresses immune sensing of self-dsRNAs through both editing-dependent (MDA5) and editing-independent (PKR, OAS-RNaseL) mechanisms. Thus, ADAR1p150 protein levels and activity combine to set the threshold for tolerance to self-derived dsRNAs.

## Introduction

Adenosine deaminase acting on RNA 1 (ADAR1) is an RNA binding and editing enzyme ^1^. ADAR1 converts adenosine nucleotides to inosine (A-to-I) in regions of double stranded RNA (dsRNA). A-to-I editing results in a permanent change to the sequence of the RNA from that encoded in the DNA. A-to-I editing is a prevalent feature of the transcriptome and can result in diverse consequences for the transcript, from changing the protein coding sequence through to modifying interactions with microRNAs and changing the secondary structure of the RNA ^2^. Most A-to-I editing in mammals occurs in repetitive elements, such as *Alu* elements (humans/primates) and short interspersed nuclear elements (SINEs, rodents) ^3, 4^. The physiologically most important function of ADAR1 is A-to-I RNA editing of cellular derived dsRNA ^1, 2, 5, 6, 7, 8, 9, 10^. This serves as a critical immune checkpoint and “self” vs “non-self” discrimination mechanism in both mouse and human by marking cellular derived dsRNA, via A-to-I editing, as self-derived and therefore not ligands for the cytoplasmic dsRNA sensor melanoma differentiation-associated protein 5 (MDA5) ^1, 5, 6, 7, 11, 12, 13^. This immune tolerance mechanism relies on the cytoplasmic ADAR1p150 isoform; the nuclear ADAR1p110 does not regulate innate sensing ^5, 14, 15, 16, 17^. Multiple mouse models including *Adar1*-null^18, 19^, isoform-specific knockouts ^14, 16, 20^, and editing-deficient mutants ^5^ demonstrate that ADAR1p150’s RNA editing activity suppresses MDA5 activation ^5, 17^.

In mammals A-to-I editing is the physiologically most important function of ADAR1p150 *in vivo*. However, editing independent, protein dependent functions for ADAR1 have been proposed ^21, 22, 23^. These include regulating RNA splicing and through protein-protein interactions that regulate microRNA formation. Although ADAR1 has been proposed to regulate microRNA biogenesis independent of its editing activity via a direct interaction with DICER, genetic analyses suggest minimal physiological relevance ^22, 24, 25, 26, 27^. Additional proposed editing independent functions such as a role for ADAR1 in senescence require further *in vivo* evaluation, now possible with viable rescued mouse models ^15, 28, 29^. Understanding the balance between ADAR1’s editing-dependent and independent functions will be important for several reasons. Firstly, this will allow a nuanced understanding of the potential impact of pathogenic variants being described in human disease such as Aicardi-Goutières’s Syndrome ^30^. Secondly, it will facilitate a more complete appreciation of the breadth of *in vivo* functions of ADAR1. Finally, it will be critical to understand how different domains of the ADAR1 protein contribute to function(s) when designing small molecules targeting ADAR1 ^1, 31, 32, 33^.

Recently, we and others demonstrated that the activation of protein kinase R (PKR) following a loss of ADAR1 was primarily dependent on RNA binding rather than editing by ADAR1p150, demonstrating a key editing independent function of ADAR1p150 protein in modulating engagement of other cytoplasmic dsRNA binding proteins ^15, 29^. Building on prior evidence of ADAR1’s potential dual functions, we employed forward genetics to dissect isoform-specific and editing-independent functions of ADAR1 *in vivo* ^15^. These analyses revealed a previously unrecognised critical role for RNA binding, but not editing, specifically by ADAR1p150 in multiple aspects of hematopoietic homeostasis *in vivo* that are independent of MDA5 and PKR. We demonstrate that ADAR1p150 is critical *in vivo* for mature peripheral T cell survival and for normal differentiation of adult hematopoietic stem and progenitor cells (HSPCs) by preventing toxicity due to sensing immunogenic self-derived RNA species formed as part of the type I Interferon-induced transcriptome. Purified ADAR1p150 deficient HSPCs were highly sensitive to low dose type I Interferon treatment *in vitro*, independent of MDA5 and PKR, and the phenotypes were ameliorated by *in vivo* antibody-mediated blockade of IFNAR1. This phenotype is conserved across multiple hematopoietic cell types. RNA binding by ADAR1p150 prevents activation of OAS-RNaseL by the cell’s own RNA induced by type I Interferon. These data demonstrate that a key editing independent function of ADAR1p150 is to sequester potentially immunogenic cellular dsRNA away from cytoplasmic dsRNA sensors, ensuring tolerance to self. Furthermore, it demonstrates that the tonic type I Interferon induced transcriptome is itself a source of cellular (‘self”) derived immunogenic dsRNAs. Therefore, ADAR1p150 is essential to tolerise the cell to self-dsRNA via both editing dependent, for MDA5, and RNA binding dependent, editing independent activities in the case of PKR and OAS-RNaseL.

## Results

### Loss of mature T cells in the peripheral blood of *Adar1^-/-^* and *Adar1^p150-/-^* triple mutants

We recently described a series of viable adult *Adar1* mutant mouse lines established by the co-deletion of MDA5 (*Ifih1^-/-^*) and PKR (*Eif2ak2^-/-^*) ^15^. These are all *Ifih1^-/-^Eif2ak2^-/-^* and are either ADAR1 wild-type and retain editing and protein expression of both ADAR1p110 and ADAR1p150 (*Adar1^+/+^*, *A1^+/+^*), are deficient in both ADAR1p110 and ADAR1p150 proteins (*Adar1^-/-^*; *A1^-/-^*) ^18^, express editing dead ADAR1p110 and ADAR1p150 proteins (*Adar1^E861A/E861A^*; *A1^EA/EA^*) ^5^ or are deficient in only the ADAR1p150 isoform (*Adar1^L196C/L196C^*; *A1^p150-/-^*)^14, 15^. This presented an opportunity to apply forward genetics to define isoform specific, editing dependent and protein dependent, editing independent functions of ADAR1 *in vivo* (**Figure 1A**).

**Figure 1:**
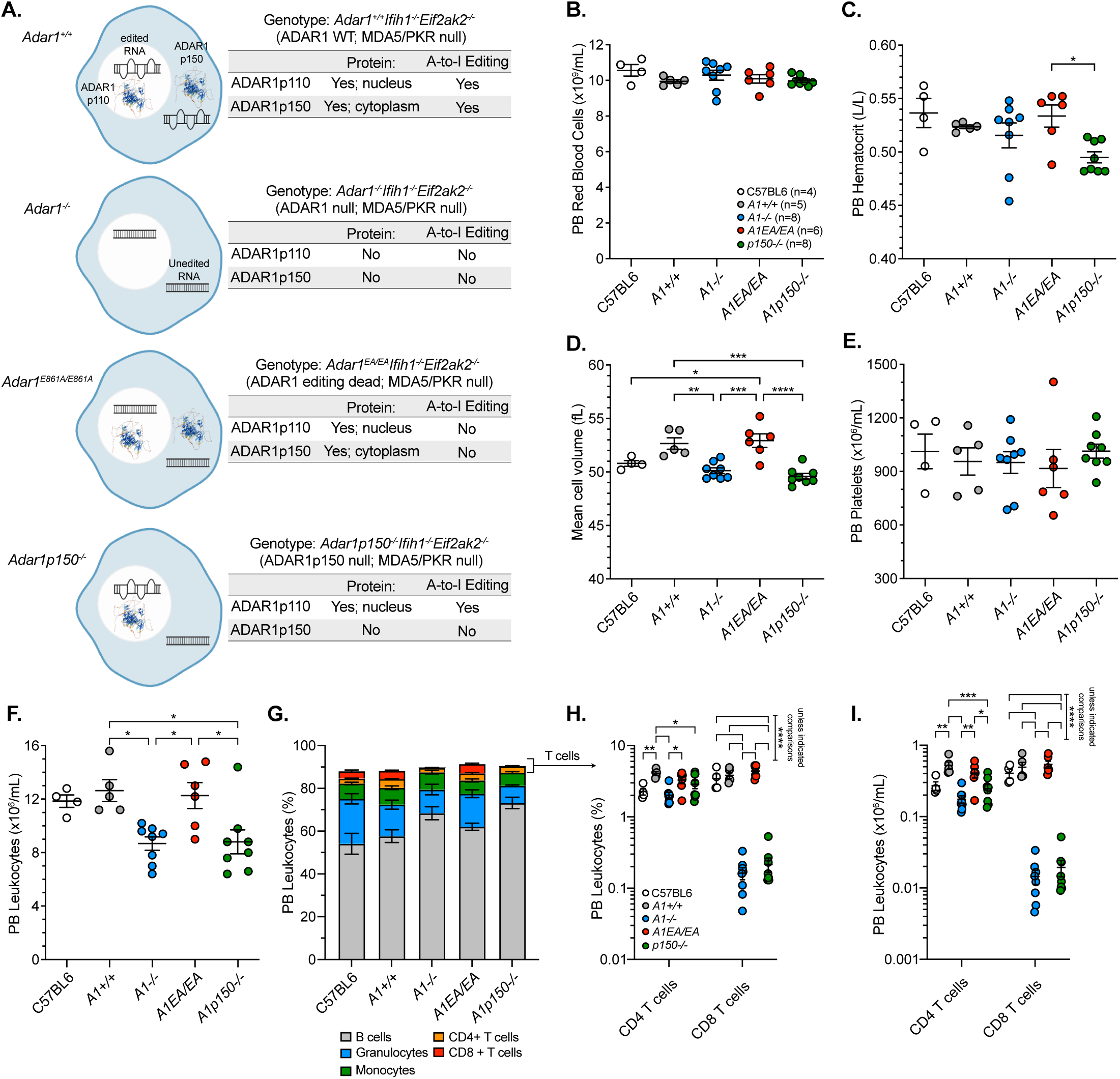
ADAR1 protein deficiency leads to a loss of peripheral blood CD4 and CD8 T cells. (**A**) Schematic outline of the ADAR1 protein status and editing status of the 4 genotypes assessed in this study. Peripheral blood (**B**) red blood cell numbers. Each data point represents an individual animal. Number of individuals per genotype as indicated (n); (**C**) Haematocrit; (**D**) Mean cell volume; (**E**) Platelet numbers; (**F**) total leukocyte numbers; (**G**) Lineage distribution of the leukocyte population; (**H**) the frequency of CD4 and CD8 T cells in each genotype; (**I**) the absolute number of CD4 and CD8 T cells per mL of PB in each genotype. Statistics calculated in Prism using a 2-way ANOVA with Tukey’s multiple comparison test correction; *p<0.05; **p<0.01; ***p<0.001, ****p<0.0001. Peripheral blood data: both males and females in dataset; Mean ± SEM as indicated. Age range of animals 62-149 days of age. Peripheral blood analysis of adult mice (age range 62-149 days). C57BL6 are CD45.1/CD45.2 compound heterozygous males (100 days of age) for reference.

Upon assessed cohorts of mixed sex adult animals from all lines with littermate *Adar1^+/+^* (*A1^+/+^*) controls we identified differences in the peripheral blood (PB) indices. There was a normal red blood cell count but a genotype dependent effect on hematocrit and mean red cell volume (**Figure 1B-1D**). The mean cell volume (MCV) in the *A1^+/+^* and *A1^EA/EA^*, both being MDA5 and PKR deficient, was increased compared to a wild-type (C57BL/6) and we had previously observed this to be MDA5 dependent ^26^. An elevated MCV was absent in the protein-deficient *A1^-/-^* and *A1^p150-/-^* animals, whose values resembled wild-type controls (**Figure 1D**). Platelets were normal (**Figure 1E**). In addition to the difference in MCV, there was a reduced PB leukocyte count in the protein deficient *A1^-/-^* and *A1^p150-/-^* cohorts but not in the editing deficient *A1^EA/EA^* animals (**Figure 1F**). Analysis of the lineage distribution of the PB leukocytes demonstrated that the most significant difference was a reduction in both the frequency and absolute number of T cells, with a ∼50% reduction in CD4 T cells and a ∼95% reduction in CD8 T cells in the PB (**Figure 1G-1I**). The other lineages in the PB were largely comparable across the genotypes demonstrating that the reduced PB T cell phenotype was due to an ADAR1 protein dependent, editing independent function (**Figure 1G**). Analyses of both murine and human single cell expression datasets indicated that ADAR1 is the most highly expressed ADAR enzyme in T cells, and hematopoietic cells more broadly (**Supplemental Figure S1A-1E**). Given that *A1^-/-^* and *A1^p150-/-^*cohorts had the same phenotypes, we could refine this to being an *in vivo* ADAR1p150 isoform specific protein dependent, editing independent function.

The T cell phenotype was dependent on ADAR1p150 protein expression, but not editing activity, so we assessed if it was dependent on the p150 specific Zα domain ^34^. The Zα domain is in the unique N-terminus of ADAR1p150 that is not shared with ADAR1p110. We assessed PB leukocytes in a Zα mutant (p.P195A) that models the most common human *ADAR* mutation pattern when combined with a null allele ^34^. *Adar1^P195A/-^* and *Adar1^P195A/+^* (control littermates; all were *Ifih1* and *Eif2ak2* wild-type) were assessed and there were no differences in the numbers of PB CD4 or CD8 cells (**Supplemental Figure S2A-2L**). This is consistent with our previous reported analysis of these lines and of the p.P195A mutants following acute somatic mutation ^34^. This suggests that the PB T cell phenotype is due to a protein dependent function of ADAR1p150 and is Zα domain and editing independent.

### Mature peripheral T cells, not developing T cells in the thymus, are absent

To more fully characterise what stage in T cell development was dependent on ADAR1p150 protein, we assessed the T cell development trajectory across a cohort of age matched, co-housed male mutants and littermate *A1^+/+^* controls (**Figure 2; Supplemental Figure S3**). The mature CD4 and CD8 T cell reductions were confirmed in the PB and the spleen (**Figure 2A-2F**). We then assessed T cell development in the thymus (**Figure 2G-2O**). Thymus cellularity was comparable across all genotypes, while thymus weight as a proportion of body weight was increased in the *A1^-/-^* only (**Figure 2G**). The proportion and absolute number of CD4+CD8+ cells in the thymus was subtlety, but significantly, different across the genotypes but the single positive CD4 or CD8 cells in the thymus were not different (**Figure 2H-2J**). There were differences in the frequency but not number of T-cell receptor (TCRβ) high CD8 single positive cells, indicative of fully mature T cells, between the *A1^EA/EA^*and *A1^p150-/-^*animals, but the *A1^p150-/-^*was not different to *A1^+/+^* (**Figure 2K-2M**). There were increased numbers of TCRβ high CD4 single positive T cells in all ADAR1 mutants compared to *A1^+/+^* (**Figure 2K-2M**). When assessing T cell development, we did not find any differences in the developing double negative (DN) stages DN1 and DN2 by either frequency or absolute number (**Figure 2N-2O**). We did find that the *A1^-/-^* had an increased proportion of DN3 compared to the *A1^p150-/-^*and that the *A1^p150-/-^*had an increased frequency of DN4. By numbers per thymus however, the DN1-DN4 cell populations were not different across all genotypes (**Figure 2O**). We also assessed hematopoiesis in the bone marrow and spleen of these cohorts, and whilst there were some differences apparent when considering population frequency, the absolute numbers of each population were largely comparable (**Supplemental Figure S3A-S3J**). Collectively these analyses indicate that, rather than a developmental block or delay in T cell development, it is most likely mature T cell survival after egress from the thymus caused the phenotypes observed in the *A1^-/-^*and *A1^p150-/-^*animals. These findings indicate that ADAR1p150 supports peripheral T cell survival through a protein-dependent, editing-independent mechanism.

**Figure 2:**
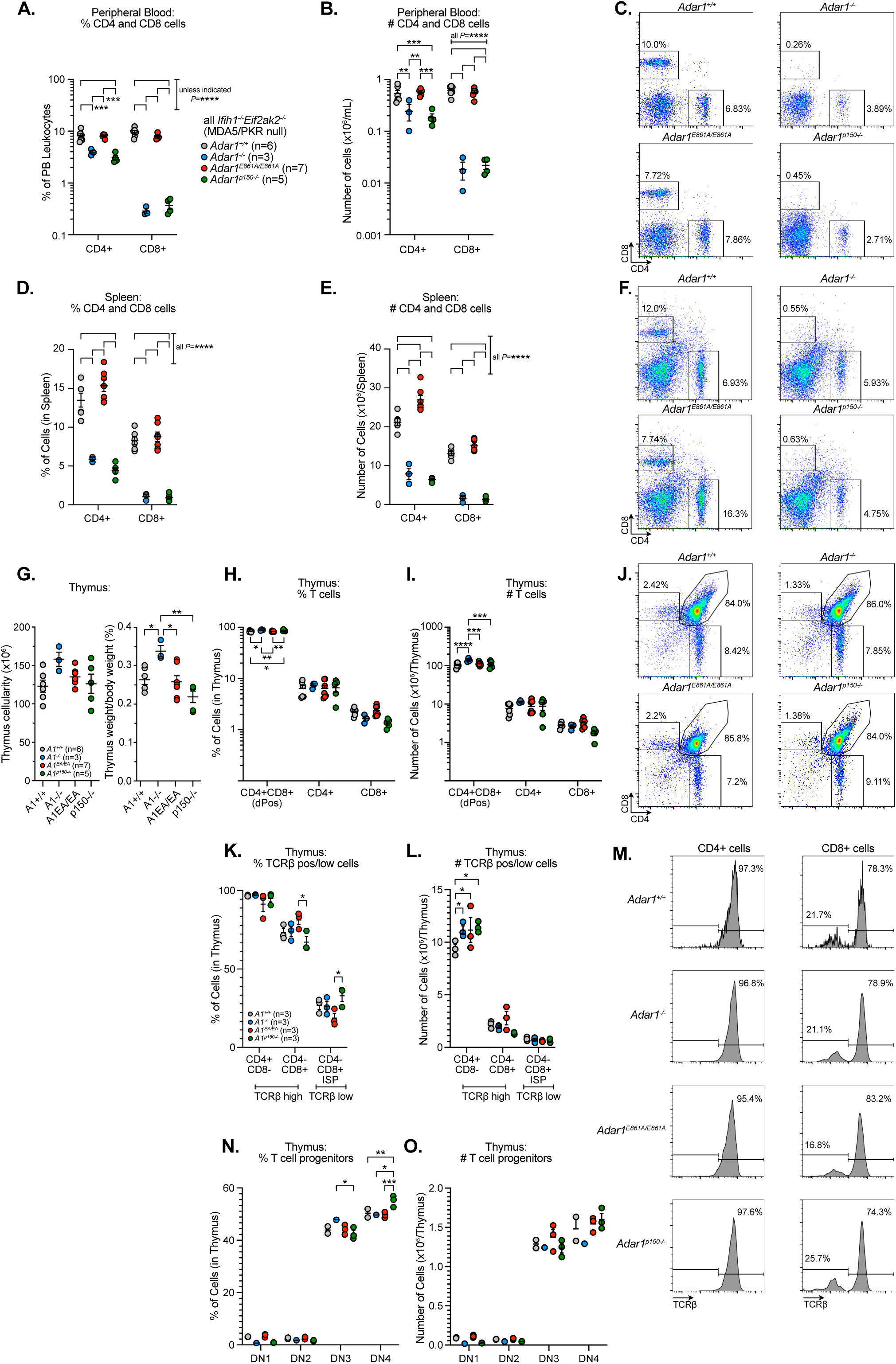
Peripheral CD4 and CD8 T cells are depleted but thymic development is largely normal. Cohorts of 10–11-week-old male animals that were littermates and co-housed were assessed. Peripheral blood T cells: (**A**) percentage of leukocytes or (**B**) number per mL; (**C**) Representative FACS plots of PB cells. Each data point represents an individual animal. Number of individuals per genotype as indicated (n). Splenic T cells: (**D**) percentage of spleen cells or (**E**) number per spleen; (**F**) Representative FACS plots of splenic cells. (**G**) Thymus cellularity and thymus weight as a proportion of body weight; (**H**) Percent of CD4+CD8+ (double positive, dPos), and single CD4+ and single CD8+ in the thymus; (**I**) Number of CD4+CD8+ (double positive, dPos), and single CD4+ and single CD8+ in the thymus; (**J**) Representative FACS plots of thymus cells stained with CD4 and CD8; (**K**) The proportion of cells of the indicated cell surface phenotype that are TCRβ high or low; (**L**) The number of cells of the indicated cell surface phenotype that are TCRβ high or low; (**M**) Representative FACS plots of TCRβ profiles of CD4 and CD8 single positive cells in the thymus; (**N**) Frequency of double negative (DN) 1 through DN4 populations in the CD4-CD8-fraction of the thymus; (**O**) Number of double negative (DN) 1 through DN4 populations in the CD4-CD8-fraction of the thymus. Mean ± SEM as indicated. Statistics calculated in Prism using a 2-way ANOVA with Tukey’s multiple comparison test correction; *p<0.05; **p<0.01; ***p<0.001, ****p<0.0001.

### Cell intrinsic defect in *Adar1^-/-^*and *Adar1^p150-/-^* triple mutant CD8 T cell production and hematopoietic reconstitution after bone marrow transplantation

Having established the peripheral T cell defect, we next asked if this was intrinsic to the hematopoietic cells or influenced by the microenvironment (both thymic and peripheral). To do so we undertook a non-competitive bone marrow transplant (BMT; **Figure 3A**) ^35^. We transplanted 5 x 10^6^ whole bone marrow from both male and female donors separately for each genotype into lethally irradiated congenic recipients. At this cell dose we would expect all recipients to have >90% chimerism from the donor cells. Analysis at 5-, 12- and 20-weeks post-transplant demonstrated that both the *A1^+/+^* and editing deficient *A1^EA/EA^* repopulated as expected and achieved >90% chimerism in the PB and normal lineage distribution of leukocytes (**Figure 3B-3M; Supplemental Figure S4A-S4P**). In contrast, the *A1^-/-^* and *A1^p150-/-^* had both lower PB leukocytes overall and significantly lower levels of PB chimerism at all time points (**Figure 3B-3M; Supplemental Figure S4A-S4P**). The poor repopulation by the *A1^-/-^* and *A1^p150-/-^* was unexpected, as when assessed under the setting of native hematopoiesis there was no indication of any defect aside from the lack of PB T cells (**Figure 1**, **Figure 2** and **Supplemental Figure S3**). The *A1^-/-^* and *A1^p150-/-^* animals that had been aged did not develop bone marrow failure ^15^. Analysis of the PB lineage distribution from the donor cells demonstrated that the *A1^-/-^* and *A1^p150-/-^* both failed to generate CD8 T cells in the wild-type recipients, demonstrating that the CD8 T cell defect was due to a cell intrinsic requirement for ADAR1p150 protein (**Figure 3G-3I; Supplemental Figure S4A-S4P**). This analysis also uncovered a protein dependent, editing independent requirement for ADAR1p150 in hematopoietic reconstitution following BMT.

**Figure 3:**
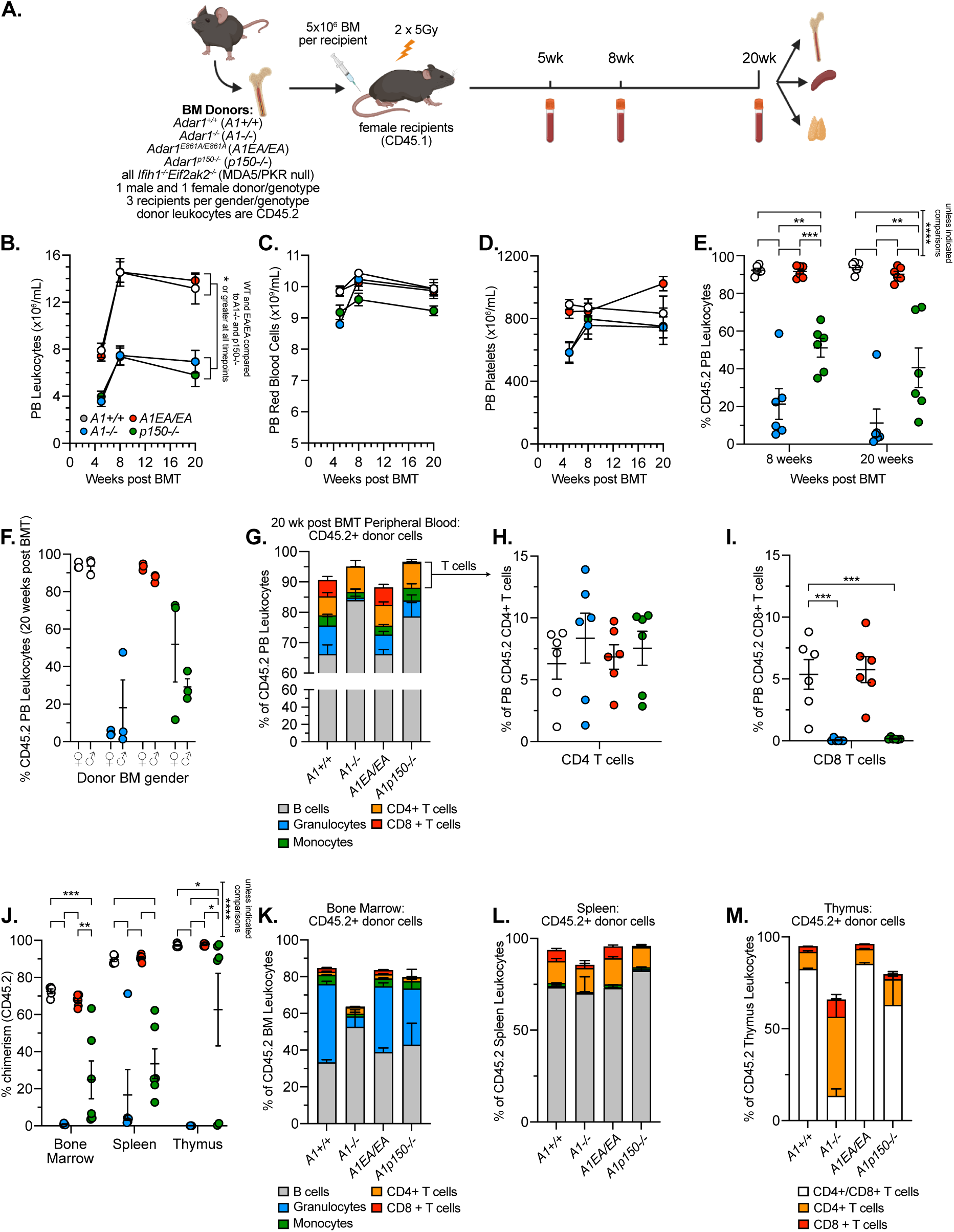
The T cell defect is hematopoietic cell intrinsic and bone marrow transplantation reveals a function of ADAR1p150 in repopulating activity. (**A**) Schematic outline of the non-competitive bone marrow transplant experiment. Peripheral blood (**B**) leukocyte number, 2-way ANOVA with Tukey’s multiple comparison test correction; compared at each timepoint, (**C**) red blood cell number, (**D**) platelet number and (**E**) percentage CD45.2 (donor) cell chimerism of leukocytes in the PB at 8 and 20 weeks post-transplant, 2-way ANOVA with Tukey’s multiple comparison test correction; compared at each timepoint; (**F**) percentage CD45.2 (donor) cell chimerism of leukocytes in the PB at 20 weeks post-transplant based on the donor cell sex; (**G**) Lineage distribution within the donor derived CD45.2+ leukocytes in the PB at 20 weeks post-transplant; (**H**) the CD4 cell and (**I**) CD8 T cells derived from donor cells at 20 weeks post-transplant, panel H and I compared using ordinary one-way ANOVA with Dunnett’s multiple comparison test; (**J**) percentage of CD45.2 cells within the leukocyte populations of the bone marrow, spleen and thymus, compared using a Mixed-effects analysis with Tukey’s multiple comparisons test; and the lineage distribution of the CD45.2 population within the (**K**) bone marrow, (**L**) spleen and (**M**) thymus. Mean ± SEM as indicated. Statistics calculated in Prism.

### Competitive repopulating activity is dependent on ADAR1p150 protein, independent of A-to-I editing activity

To determine whether the observed repopulating defect was a reflection of impaired stem cell activity, we performed a competitive BMT. In this setting, the *A1* mutant cells (CD45.2) were mixed 1:1 with CD45.1/CD45.2 heterozygous congenic bone marrow and transplanted into CD45.1 lethally irradiated recipients (**Figure 4A**). This provides a common internal CD45.1/CD45.2 control population in each recipient and across each genotype as a comparator. Analysis of PB was completed at 5-, 12- and 20-weeks post-transplant. While the *A1^+/+^*and editing-deficient *A1^EA/EA^* showed robust repopulation, the *A1^-/-^* and *A1^p150-/-^*had profoundly reduced competitive repopulating activity and both had significantly reduced chimerism across all lineages, indicative of a significant defect at the hematopoietic stem cell (HSC) or primitive progenitor level (**Figure 4B-4G; Supplemental Figure S5A-S5F**). We quantitated the competitive repopulating units (CRU), an estimate of HSC frequency ^35, 36, 37, 38^. When compared to the wild-type competitor cells (wild-type for *Adar1*, *Ifih1* and *Eif2ak2*), which have a CRU activity of 10 based on the 1×10^6^ whole bone marrow cells used, the *A1^+/+^*had 2.8-fold increased CRU numbers at 20 weeks post-transplant (**Figure 4E**). The editing deficient *A1^EA/EA^* had a CRU number that was the same as the wild-type competitor cells (**Figure 4E**). In contrast, the protein deficient *A1^-/-^* and *A1^p150-/-^* had a ∼600-fold reduction compared to the *A1^+/+^* and ∼200-fold reduced CRU compared to the editing deficient *A1^EA/EA^* (**Figure 4E**). We independently replicated these findings with the *A1^p150-/-^* deficient cells and *A1^+/+^*controls (**Figure S6A-S6D**). These studies reveal several unanticipated findings. Firstly, the absence of MDA5 and PKR resulted in improved competitive repopulating activity, evidenced by the increased CRU and chimerism of the *Adar1^+/+^Ifih1^-/-^Eif2ak2^-/-^* cells. Interestingly, loss of editing in the *A1^EA/EA^* cells negated this advantage. Secondly, ADAR1p150 protein is required for both non-competitive and competitive HSC repopulation following BMT independent of its editing activity. Previously, studies had identified a critical function for ADAR1 in HSC homeostasis, but it was not clear if there was a contribution independent of editing or MDA5 ^5, 19^. These analyses demonstrate a new function for ADAR1p150 protein in HSC repopulating potential.

**Figure 4:**
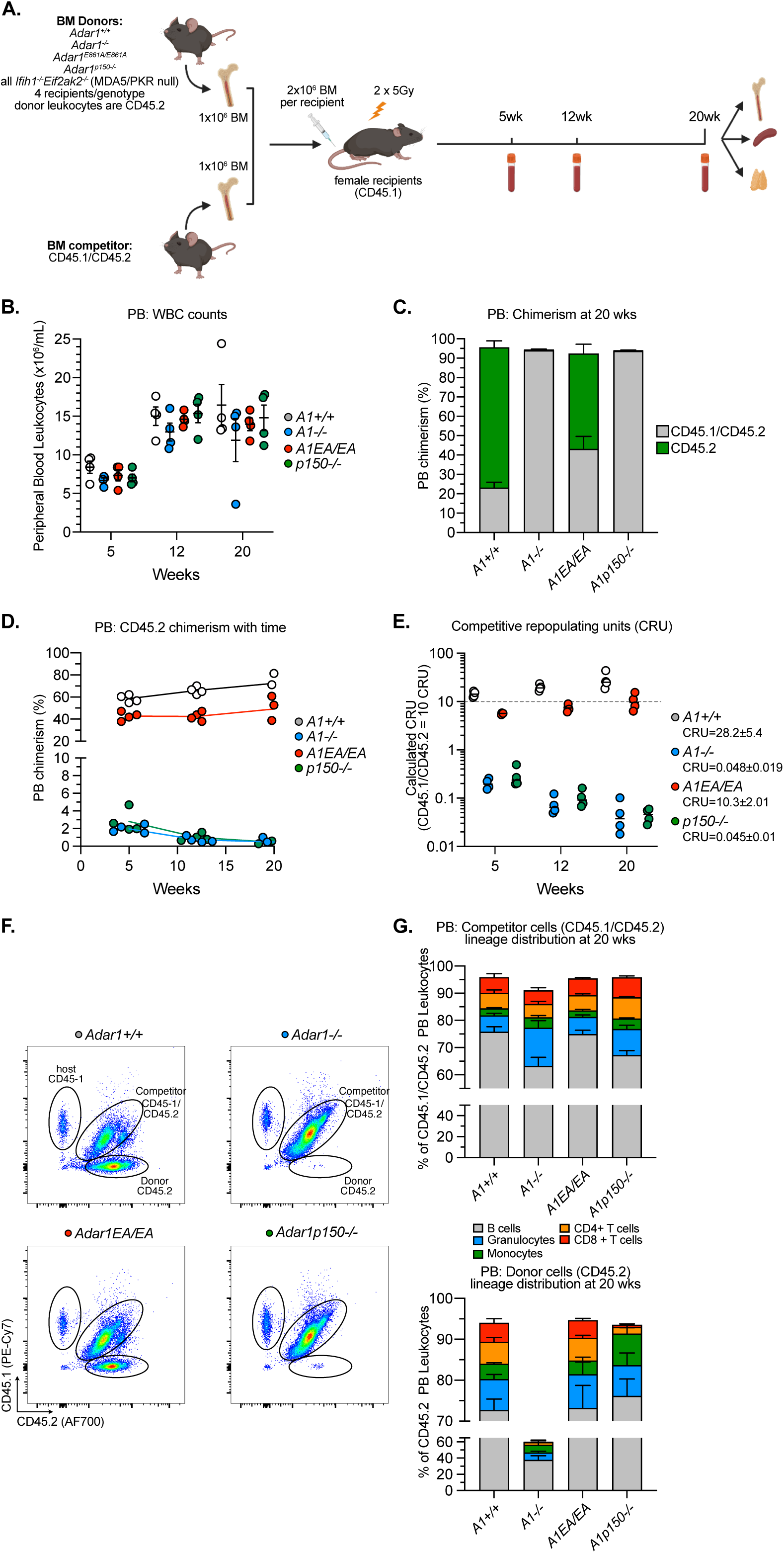
Severely reduced competitive repopulating potential dependent on ADAR1p150 protein but not editing. (**A**) Schematic outline of the competitive bone marrow transplant experiment. Peripheral blood (**B**) leukocyte number and (**C**) percentage donor (CD45.2) compared to competitor (CD45.1/CD45.2) chimerism in the leukocyte population at 20 weeks post-transplant. (**D**) Donor cell CD45.2 chimerism in the PB across time by genotype. (**E**) Calculated competitive repopulating units (CRU) at each time point assessed; The competitor BM (CD45.1/CD45.2) is assigned a CRU of 10 based on 1 CRU per 1x 10^5^ bone marrow leukocytes; the dashed bar indicates the CRU=10 level; calculated CRU per genotype as indicated. (**F**) Representative FACS plots of CD45.1 vs CD45.2 stained PB leukocytes and (**G**) lineage distribution within the competitor (upper panel) or donor (lower panel) cells. Where applicable each data point represents as individual animal or data expressed as mean ± SEM, n=4 recipients per genotype.

### Hypersensitivity to type I Interferon underlies both the HSC and T cell defects

In the setting of native hematopoiesis there was a restricted phenotype impacting the mature CD4 and CD8 T cells in the PB, whilst following the stress of BMT we observed a profound HSC repopulating defect. To understand if these phenotypes were molecularly linked or separable, we considered the context of each setting. Type I interferon (IFN) has been demonstrated to be important in survival and activation of mature T cells and is produced by thymic epithelial cells in sterile conditions ^39, 40, 41^. When exposed to total body irradiation, cell death induced in the host cells results in the significant release of inflammatory cytokines including type I IFNs ^42, 43, 44^. HSCs can be directly impacted by type I IFN and chronic type I IFN exposure leads to depletion of HSCs and compromises their function and transplant activity ^45, 46, 47, 48, 49, 50, 51, 52^.

To directly test the response of the *A1^p150-/-^* deficient HSPCs to type I IFN, we isolated HSCs and multiple primitive cell populations from *A1^p150-/-^* and littermate *A1^+/+^* controls and treated these *in vitro* with recombinant murine IFNβ (rmIFNβ). During purification of the cell populations from animals in the context of native hematopoiesis, we observed that the *A1^p150-/-^* HSPCs had an altered differentiation trajectory compared to the *A1^+/+^* controls. In *A1^+/+^*controls the HSC population transitioned to a phenotypic ST-HSC population then progressed through the multipotent progenitor 3 (MPP3) to MPP4 stages ^53^ (**Figure 5A**). In contrast, the *A1^p150-/-^* HSCs did not transition through the phenotypic ST-HSC fraction, transitioning via an HSC to MPP2 to MPP3 route instead (**Figure 5B**). This resulted in an ∼80% reduction in the frequency of ST-HSCs and an expansion of the MPP2 population (**Figure 5C**). It is noteworthy that the altered differentiation trajectory of the *A1^p150-/-^* HSPCs does not result in profound defects in hematopoiesis, except for the described CD4/CD8 T cell defect in the PB, throughout life (**Figure 1**, **Figure 2, Supplemental Figure S3**) ^15^. We purified the phenotypic LT-HSC, MPP1, ST-HSC and MPP4 populations ^54^, treated these cells with 50U/mL rmIFNβ in serum free culture conditions for 72 hours and assessed the viability (**Figure 5D-5E**). Sufficient MPP4 were recovered to also treat with 100U/mL rmIFNβ. The *A1^p150-/-^* ST-HSCs were more sensitive to low dose rmIFNβ, with a significant reduction in total cell number and viability compared to *A1^+/+^* ST-HSCs (**Figure 5D-5E**). This phenotyping and *in vitro* analysis demonstrated that the ADAR1p150 protein, independent of its RNA editing activity, was essential *in vivo* for normal HSPC differentiation and for buffering the response to sterile type I IFN exposure *in vitro*.

**Figure 5:**
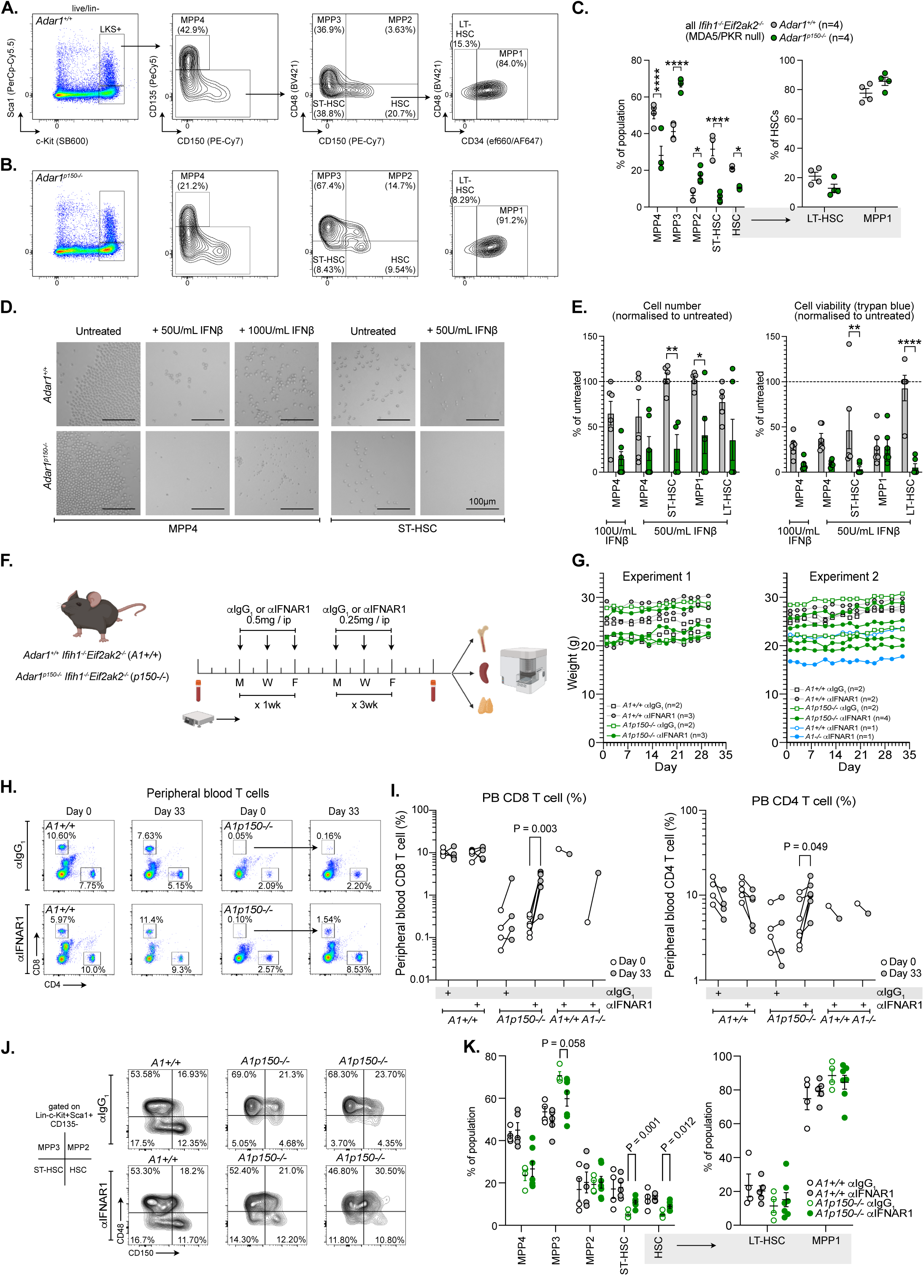
ADAR1p150 protein deficiency results in type I IFN hypersensitivity *in vitro* and blockade of IFNAR *in vivo* rescues the T cell and HSPC phenotypes. (**A**) Representative FACS plots from an *Adar1^+/+^Ifih1^-/-^Eif2ak2^-/-^* (*A1^+/+^*) showing the gating strategy to identify the HSC and progenitor fraction. (**B**) Representative FACS plots from an *Adar1^p150-/-^ Ifih1^-/-^Eif2ak2^-/-^*(*A1^p150-/-^*) animal. Experiment repeated 4 independent times with 4 pairs of control and *A1^p150-/-^* males each time. (**C**) Frequency of each population in each genotype; each data point represents as individual animal and mean ± SEM as indicated, n=4 per genotype. (**D**) Representative microscope images for each indicated cell population in each treatment arm. (**E**) The cell number for each population expressed as a percentage of untreated cells (left panel) and viability as determined by trypan blue cell counting expressed as a percentage of untreated cells (right panel); data expressed as mean ± SEM, n=4 per genotype. (**F**) Schematic outline of *in vivo* experiment to block IFNAR1 using monoclonal antibody administration. (**G**) Body weight of each animal administered αIgG_1_ or αIFNAR1 monoclonal antibody in each experiment. Each experiment was performed with an independent batch of αIFNAR1 with n per genotype as indicated. (**H**) Representative FACS plots of peripheral blood T cell populations. (**I**) Quantitation of the peripheral blood CD8 (left panel) and CD4 (right panel) T cells in each genotype and treatment cohort; Data from each individual animal is linked by a line and all animals across both replicates pooled. Statistical significance determined by multiple paired T test with FDR correction for multiple comparisons. (**J**) Representative FACS plots of bone marrow HSPC populations; populations gated on lineage^-^ cKit^+^ Sca1^+^ CD135^-^ population. (K) Frequency of each indicated population in each genotype and treatment cohort; each data point represents as individual animal and mean ± SEM as indicated and all animals across both replicates pooled, Statistical significance determined by multiple paired t-tests with FDR correction for multiple comparisons.

Given the result with purified HSPCs and known role of type I IFN in T cell biology, we directly tested if blocking IFNAR1 could rescue the peripheral T cell defect in the *A1^p150-/-^* mice *in vivo*. *A1^p150-/-^* and littermate *A1^+/+^* control mice were treated with an α-IFNAR1 neutralising monoclonal antibody or α-IgG_1_ isotype control for 4 weeks (**Figure 5F**) ^55, 56^. A single *A1^-/-^* and control *A1^+/+^* littermate were also treated with the α-IFNAR1 neutralising antibody. Peripheral blood was assessed prior to treatment and after 33 days of treatment. All genotypes tolerated the treatment well (**Figure 5G; Supplemental Figure S7A-S7G**). After 4 weeks treatment with the α-IFNAR1 neutralising antibody both the CD8 and CD4 T cell frequency and numbers in the PB of the *A1^p150-/-^* animals significantly increased (**Figure 5H-5I; Supplemental Figure S7A-S7D**). Rescue also occurred for CD8 T cells in the PB of the single *A1^-/-^* assessed. At completion of treatment, we assessed the bone marrow HSPC fractions. Strikingly, blockade of endogenously derived tonic type I IFN *in vivo* re-established the HSPC differentiation trajectory and resulted in a recovery of the ST-HSC phenotypic population in the *A1^p150-/-^* animals (**Figure 5J-5K; Supplemental Figure S7H-S7I**). Collectively, these results demonstrate that the reduced CD4 and CD8 cells in the PB and altered HSPC differentiation trajectory of the *A1^p150-/-^* animals is due to hypersensitivity to physiologic sterile type I IFN signaling. This also indicates that type I IFN hypersensitivity is a generalisable feature of ADAR1p150 protein deficient hematopoietic cell types and not restricted to a single lineage.

### RNaseL activation mediates cell death in *Adar1^-/-^* and *Adar1^p150-/-^*triple mutant myeloid cell lines following low dose type I Interferon

To understand how the loss of ADAR1p150 protein was causing hypersensitivity to type I IFN we sought a tractable cell line model. We generated HoxA9 immortalised myeloid progenitor cells from the four *Adar1* genotypes as we have previously reported (3 cell lines per genotype, each cell line derived from a separate bone marrow donor; **Figure 6A**) ^15, 34, 57^. We first determined if the immortalised myeloid cell lines were hypersensitive to type I IFN. Treatment with a low dose (50U/mL) or high dose (500U/mL) rmIFNβ resulted in the death of both the *A1^-/-^* and *A1^p150-/-^* but not the *A1^+/+^*or editing deficient *A1^EA/EA^* cells (**Figure 6B-6C; Supplemental Figure S8A-S8C**). The increased cell death of the ADAR1p150 protein deficient cells was apparent using both trypan blue cell counts and upon continuous monitoring using Cytotox viability dye in an Incucyte (**Figure 6B-6C; Supplemental Figure S8F-S8G**). This demonstrates that hypersensitivity to type I IFN is a common feature of ADAR1p150 protein deficient hematopoietic cell types and that this is not dependent on RNA editing, MDA5 or PKR.

**Figure 6:**
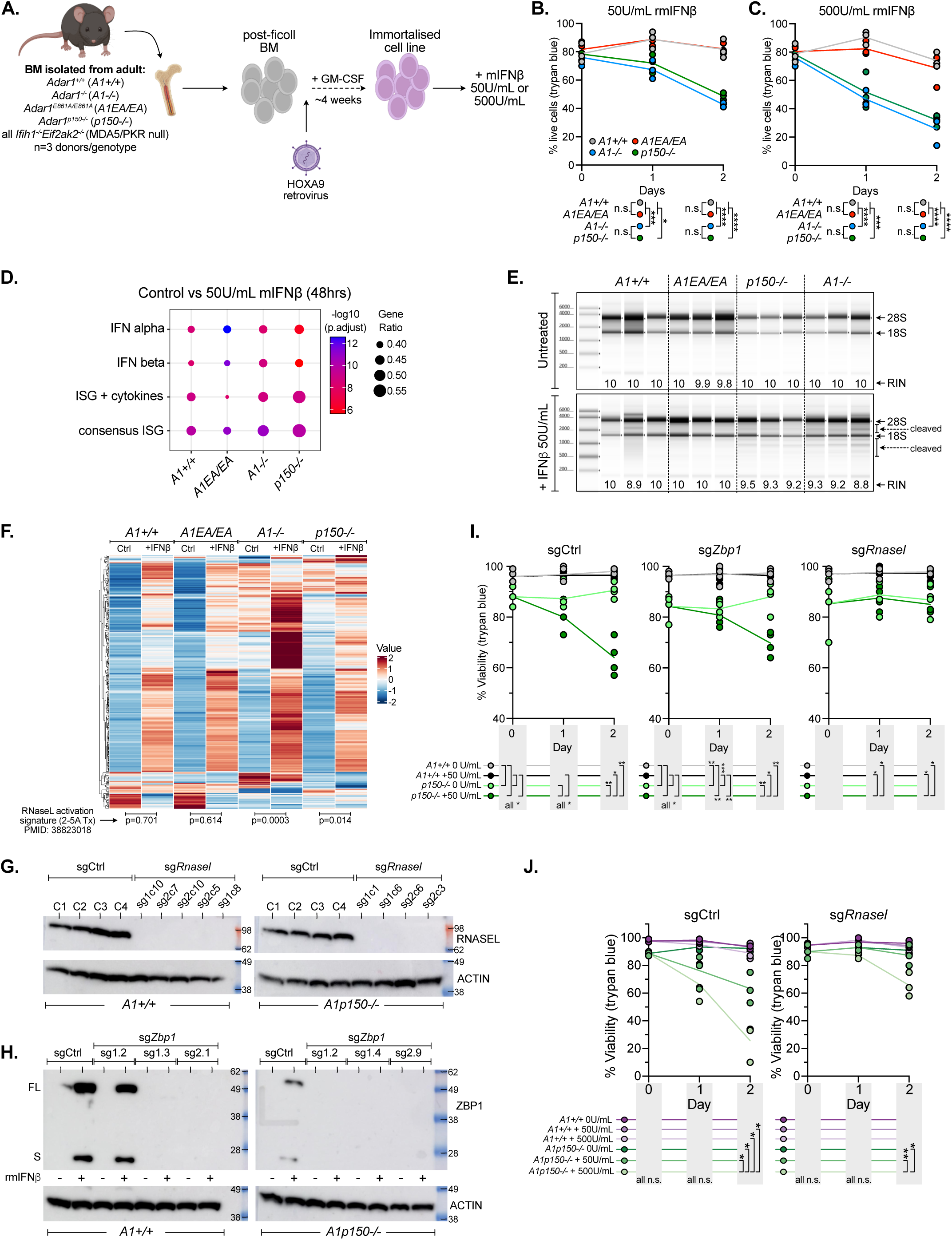
OAS-RNaseL is activated by type I IFN in the absence of ADAR1p150 protein. (**A**) Schematic outline of the generation of immortalised myeloid cell lines; n=3 independent bone marrow donors per genotype were used to establish the cell lines. (**B**) Percentage of live cells (compared to untreated time 0; trypan blue staining) following treatment with 50U/mL rmIFNβ for 48 hours; each individual cell line indicated by a dot; statistics calculated using 2-way ANOVA with Tukey’s multiple comparison correction. (**C**) Percentage of live cells (compared to untreated time 0; trypan blue staining) following treatment with 500U/mL rmIFNβ for 48 hours; each individual cell line indicated by a dot; statistics calculated using 2-way ANOVA with Tukey’s multiple comparison correction. (**D**) Analysis of indicated gene signatures in RNA-seq of untreated cells and cells treated 50U/mL rmIFNβ for 48 hours; n=3 cell lines per genotype. (**E**) RNA quality assessment of the RNA used in RNA-seq in panel D; the 28S and 18S ribosomal RNAs as indicated and cleaved RNA pattern indicated. (**F**) Analysis of enrichment of an RNaseL activation signature in the RNA-seq data from the 48 hour treated samples. (**G**) Validation of generated sgControl (sg*Ctrl*) and RNASEL deficient (sg*RnaseL*) clones in the *Adar1^+/+^Ifih1^-/-^Eif2ak2^-/-^*(*A1^+/+^*) and *Adar1^p150-/-^Ifih1^-/-^ Eif2ak2^-/-^* (*A1^p150-/-^*) cell lines. For panels G-J: A1+/+ cell line = #214; A1p150-/- cell line = #222; 3-4 CRISPR clones per gene per cell line (meaning 3 or 4 different targeting regions sgRNA per line; individual lines clones. Indicated number of clones per genotype were isolated with mutations confirmed by Sanger sequencing and western blot. (**H**) Validation of generated sgControl (sg*Ctrl*) and ZBP1 deficient (sg*Zbp1*) clones in the *Adar1^+/+^Ifih1^-/-^Eif2ak2^-/-^*(*A1^+/+^*) and *Adar1^p150-/-^Ifih1^-/-^ Eif2ak2^-/-^* (*A1^p150-/-^*) cell lines. Treatment with 50U/mL rmIFNβ for 24 hours was used to induce expression of ZBP1. Indicated number of clones per genotype were isolated with mutations confirmed by Sanger sequencing and western blot. Only clones showing loss of both short and long isoforms of ZBP1 were used for further analysis. (**I**) Treatment of the indicated genotypes with 50U/mL rmIFNβ for 48 hours and viability plotted as a percentage of untreated cells (time 0) for each line; each individual cell line indicated by a dot; statistics calculated using 2-way ANOVA with Tukey’s multiple comparison correction. (**J**) Replicate of experiment shown in panel I including dosing with 500U/mL rmIFNβ for 48 hours and viability plotted as a percentage of untreated cells (time 0) for each line; each individual cell line indicated by a dot; Mean ± SEM as indicated; Statistics calculated using 2-way ANOVA with Tukey’s multiple comparison correction.

Having established a cell line model that recapitulated the hypersensitivity to type I IFN, we sought to understand how this was occurring. Transcriptomic analyses of control and rmIFNβ treated cells both at early time points (0.5, 1, 4hrs of treatment) or after 48 hours demonstrated that the response was comparable across all genotypes (**Figure 6D; Supplemental Dataset S1 and Figure S10**). This indicated that a different magnitude of response to type I IFN between genotypes was not the likely the cause of the phenotype (**Figure 6D; Supplemental Dataset S1 and Figure S10**). The cell surface levels of IFNAR1 were the same across all genotypes (**Supplemental Figure S8D**), as were the gene expression levels of the type I IFN receptors (**Supplemental Figure S8E**). The magnitude of change in interferon stimulated genes (ISGs) was comparable across all genotypes when individual genes were assessed (**Supplemental Figure S9A**). Therefore, the absence of ADAR1p150 protein was not resulting in a fundamentally altered response. When assessing the RNA integrity of samples at 48 hours we noted the presence of banding patterns consistent with RNaseL activation in the *A1^-/-^* and *A1^p150-/-^*samples but not the *A1^+/+^* and *A1^EA/EA^* samples (**Figure 6E**) ^58^. RNaseL mediates the endonucleolytic cleavage of RNAs, both viral and ribosomal, in response to infection or interferon stimulation ^58^. RNaseL does not directly sense RNA but is activated by 2′-5′-linked oligoadenylates (2-5A) produced following cytoplasmic RNA detection by the oligoadenylate synthetase (OAS) family of RNA sensors ^59^. The transcripts encoding the OAS-RNaseL system were expressed (counts per million (CPM)>1) at baseline in all genotypes prior to rmIFNβ treatment (**Supplemental Figure S9B**). We assessed the transcriptomic datasets and there was evidence of an RNaseL activation signature in the *A1^-/-^* and *A1^p150-/-^* but not the *A1^+/+^* and *A1^EA/EA^* samples (**Figure 6F; Supplemental Figure S10A-S10B**)^60^.

Both the RNA integrity analysis and the transcriptomic signature suggested RNaseL was being activated following type I IFN signaling in the absence of ADAR1p150 protein. To confirm if this caused the cell death, we generated *A1^+/+^* and *A1^p150-/-^* myeloid cell clones that were deficient in either *RnaseL* or *Zbp1*. We assessed if ZBP1 was involved due to recent literature linking it to phenotypes associated with loss or mutation of ADAR1 ^61, 62, 63, 64, 65, 66^. As for the transcripts encoding the OAS-RNaseL system, *Zbp1* and related putative effectors such as *Ripk1/2/3/Mlkl* were all expressed (counts per million (CPM)>1) at baseline in all genotypes prior to rmIFNβ treatment and many displayed the characteristic interferon induced increase in gene expression (**Supplemental Figure S9B-S9C**). This indicated that these sensing systems are expressed but inactive in the absence of type I IFN treatment. After confirming that the cells were deficient for RNaseL or both short and long isoforms of ZBP1 (**Figure 6G-6H; Supplemental Figure S12**), we treated the cells with 50U/mL rmIFNβ for 48 hours. Strikingly, the loss of ZBP1 did not change the response, with the sg*Zbp1^-/-^ A1^p150-/-^* myeloid cell lines remaining sensitive to type I IFN (**Figure 6I**). In contrast, and consistent with the transcriptomic signature and RNA integrity analysis, the loss of RNaseL completely protected against cell death induced by low dose rmIFNβ for 48 hours (**Figure 6I-6J**). This demonstrates that ADAR1p150 protein, but not editing, prevents OAS-RNaseL activation following low dose type I IFN treatment.

### ADAR1p150 RNA binding prevents RNaseL activation by the cellular type I interferon induced transcriptome

To determine how ADAR1p150 protein suppressed OAS-RNaseL activation, we expressed murine ADAR1 variants in the myeloid cell lines and re-exposed them to rmIFNβ for 48 hours. We introduced wild-type ADAR1p150, an RNA binding mutant ADAR1p150 generated by mutating the KKxxK motif to EAxxA in all three dsRNA binding domains (dsRBD) and wild-type ADAR1p110 (**Figure 7A**) ^15, 67, 68^. ADAR1p110 is nuclear, which allowed us to understand if the critical function was exclusively performed in the cytoplasm. All cDNA were N-terminal GFP tagged and had a 3xFlag tag on the C-terminus ^68^. Cells were infected, selected and the GFP positive population purified by FACS, protein expression confirmed by western blotting and activity of the protein on a cellular transcript known to be edited (**Figure 7B**). The cells were then treated with 50U/mL or 500U/mL rmIFNβ for 48 hours. The restoration of wild-type ADAR1p150 prevented cell death following treatment, as expected (**Figure 7C**). ADAR1p110 expression did not prevent cell death, consistent with the failure of the retained endogenous ADAR1p110 in the *A1^p150-/-^* myeloid cell lines to prevent toxicity from type I IFN (**Figure 7C**). This demonstrates that cytoplasmic ADAR1 protein localisation is essential. The RNA binding ADAR1p150 mutant failed to prevent cell death following treatment with either 50U/mL or 500U/mL rmIFNβ for 48 hours (**Figure 7C**). This establishes that it is the type I IFN induced transcriptome that results in cell death, indicating that the RNAs that are induced as part of this response are themselves immunogenic and form substrates that can be sensed by OAS-RNaseL. These results establish that ADAR1p150 RNA binding, not editing, acts to prevent OAS-RNaseL activation by cellular derived RNA species following type I IFN treatment.

**Figure 7:**
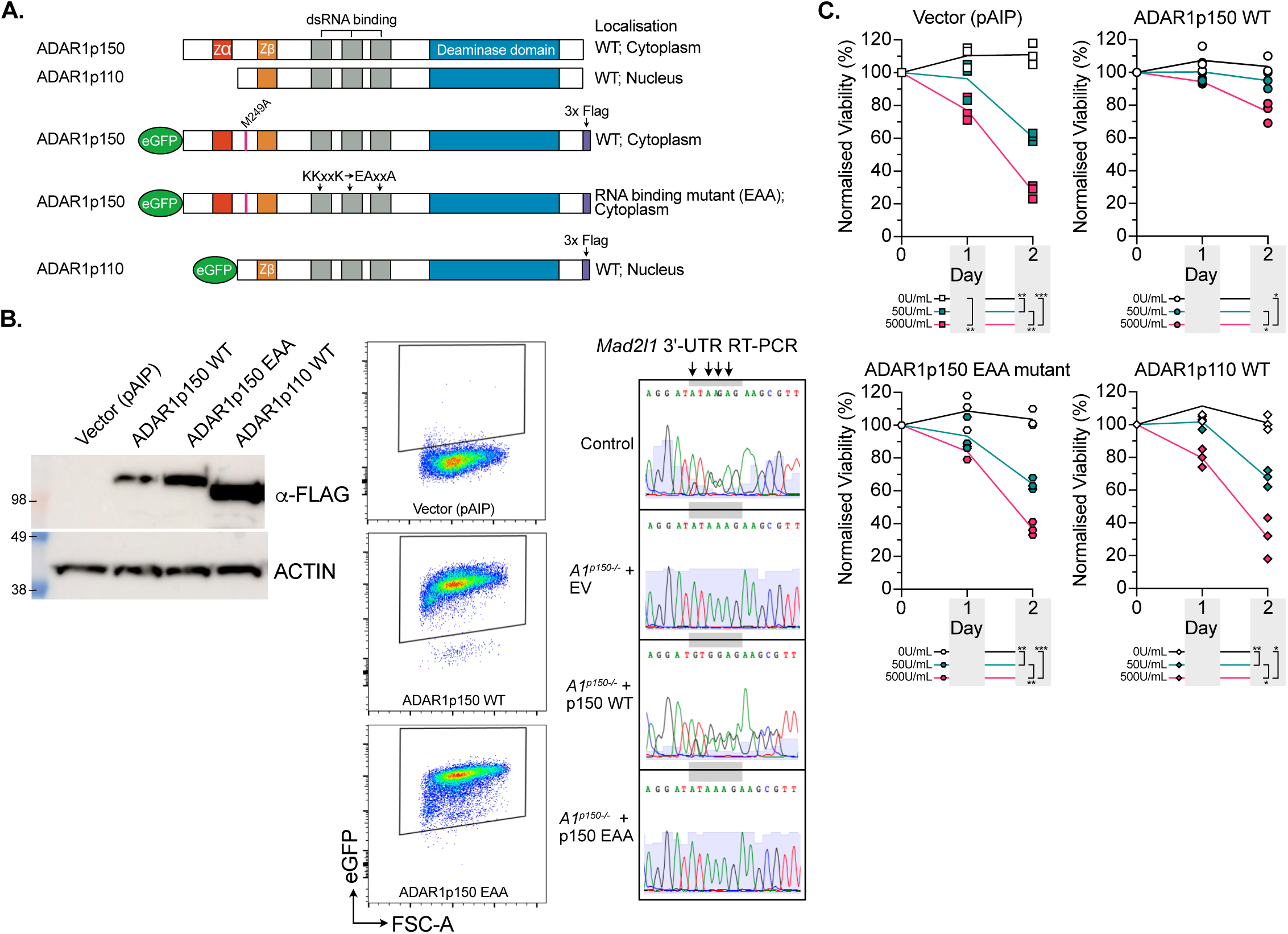
RNA binding by ADAR1p150 prevents toxicity from type I IFN treatment. (**A**) Schematic outline of the ADAR1 cDNAs used for re-expression. All cDNAs were re-expressed in the *Adar1^p150-/-^Ifih1^-/-^Eif2ak2^-/-^* (*A1^p150-/-^*) cell genotype only. (**B**) Western blot (using α-Flag) of FACS purified cells ectopically expressing ADAR1 (left), representative FACS plots of the GFP expression in each indicated genotype (middle) and Sanger sequencing traces of the *Mad2l1* 3’UTR demonstrating restoration of editing as indicated by the arrows by the WT but not EAA mutant ADAR1 (control from are *Adar1^E861A/+^* cell line). (**C**) Percentage of live cells (compared to untreated time 0; trypan blue staining) following treatment with 50U/mL or 500U/mL rmIFNβ for 48 hours; each individual cell dot represents an independently infected/FACS sorted sample of each genotype; statistics calculated using 2-way ANOVA with Tukey’s multiple comparison correction.

## Discussion

Sterile activation of innate immune dsRNA sensors by endogenous RNA is increasingly recognized as a key driver of inflammation and autoimmunity ^69, 70, 71, 72^. ADAR1-mediated A-to-I editing of dsRNA is critical for regulating immunogenicity and tolerance to self-RNA ^1, 2, 11, 14, 15, 73^. This editing, primarily by cytoplasmic ADAR1p150, prevents MDA5 from sensing under-/un-edited cellular dsRNA, a conserved mechanism across species ^1, 5, 7, 73^. MDA5 activation triggers a potent innate immune response, including type I interferon (IFN) production and cell death ^5, 18, 19^. While ADAR1’s editing role is well established, its editing-independent functions remain poorly understood. We recently showed that PKR, another cytoplasmic RNA sensor, is activated downstream of MDA5 and suppressed by ADAR1p150 protein ^15^, confirmed independently ^29^. Notably, RNA binding, not editing, by ADAR1p150 prevented PKR from engaging cellular transcripts ^15^. The broader impact of RNA binding by ADAR1 independent of A-to-I editing is unclear. Using forward genetics, we analysed an allelic series of *Adar1* mutant mice rescued by loss of both MDA5 and PKR (phenotype summarised in **Supplemental Figure S11A**). This allowed us to uncover editing-independent functions of ADAR1p150, including its role in buffering toxicity from tonic type I IFN *in vivo* ^74^, a previously unobservable interaction due to lethality in ADAR1-deficient mice.

In ADAR1p150-deficient mice, we observed significant loss of mature CD4+ and CD8+ T cells in the peripheral blood and organs. Surprisingly, thymic T cell development remained largely intact, differing from prior studies where MDA5 was still present ^75^. Bone marrow transplant experiments confirmed this was due to a T cell-intrinsic role of ADAR1p150. Our findings reveal that ADAR1p150 is crucial for maintaining mature T cells post-thymic egress, identifying it as a novel regulator of peripheral T cell survival and homeostasis. The *in vivo* source of tonic type I IFN driving mature T cell death remains unknown. Type I IFNs are physiologically produced and essential for hematopoietic cell homeostasis. For example, sterile Type I and III IFN production by AIRE+ thymic epithelial cells affects T cell homeostasis ^41^. HSCs *in vivo* can also respond to type I IFN and inflammatory stimuli that cause a type I IFN response in an IFNAR dependent manner ^76^. Although ADAR1 has been linked to T-lymphoblastoid leukemia (T-ALL), those effects were MDA5-dependent, unlike our model ^77^. Prior overexpression studies suggested both editing-dependent and independent roles for ADAR1p150 in regulating type I IFN toxicity ^77^. However, our data conclusively show that its role in T cell homeostasis is editing independent. While ADAR1p150’s editing-dependent suppression of MDA5 remains its dominant function, we can now appreciate editing-independent roles in T cell regulation. This has important implications for therapeutic targeting of ADAR1. Strategies that eliminate the protein or disrupt its RNA-binding domains may inadvertently impair T cell homeostasis, potentially weakening T cell-mediated anti-tumor immunity, a major goal of ADAR1 inhibitor development ^31, 32, 33, 78^.

We previously aged various *Adar1* mutants and did not observe hematopoietic deterioration, such as bone marrow failure ^15, 26, 27^. Thus, we were surprised to find altered HSPC differentiation following ADAR1p150 protein loss. Previously we had identified ADAR1’s role in HSC maintenance and it appeared editing-dependent, with MDA5 mediating HSPC loss ^5, 19, 26^. Our current findings extend this understanding, showing ADAR1p150 can contribute to HSC homeostasis and primitive progenitor differentiation independently of editing and MDA5/PKR. Loss of ADAR1p150 led to ∼80% reduction in ST-HSCs, with differentiation shifting to an alternative HSC to MPP2 to MPP3 pathway. Despite this, mature lineages (excluding T cells) were produced normally, indicating hematopoiesis accommodates this altered differentiation pathway under native conditions. However, ADAR1p150 protein becomes essential under stress, such as after bone marrow transplantation (BMT). Future studies should explore other stressors, such as bleeding, infection or chemotherapy, to assess ADAR1p150’s role in HSC regeneration. The ability of ADAR1p150 protein to protect HSCs from transplant stress was not possible to appreciate previously and demonstrates that ADAR1p150 has multifaceted roles in HSC biology that are both editing dependent and independent.

A unifying feature of the impaired mature peripheral T cell survival and compromised HSC transplant activity is hypersensitivity to type I IFN. In native hematopoiesis, IFN signaling is tonic; in BMT, it is induced by the irradiation used in conditioning of recipients ^42, 43, 44^. Purified HSCs and MPPs lacking ADAR1p150 protein showed increased sensitivity to low-dose IFN *in vitro*. Blocking IFNAR1 *in vivo* significantly improved peripheral blood T cell numbers and normalised HSPC differentiation, directly linking these phenotypes to type I IFN activity or response. Using immortalized myeloid lines, we confirmed type I IFN sensitivity was ADAR1p150-protein dependent, not editing-related, and a common feature across distinct types of hematopoietic cells spanning the differentiation continuum.

ADAR1p150 suppressed activation of the OAS-RNaseL pathway triggered by cellular type I IFN-induced transcripts. OAS proteins sense cytoplasmic dsRNA and produce 2’-5’-A, activating RNaseL, a mechanism normally associated with antiviral response ^59^. Whilst the preferred RNA ligands for OAS’s have been described in various settings ^79, 80, 81^, the ligands in our settings are cellular and arise from sterile type I IFN induced transcription. Here ADAR1p150 prevents OAS sensing of cellular dsRNAs in an RNA binding dependent manner. Our data indicate that the mature T cells and the HSPC populations experience tonic physiological type I IFN exposure in their normal microenvironments under homeostatic conditions. In the case of mature T cells this could be anywhere in the periphery while for HSPCs this is within the bone marrow microenvironment. The specificity of the response is intriguing, suggesting either differential thresholds for sensitivity across affected compared to unaffected cell types or highly localised and focal type I IFN production and signalling that drives the cellular specificity *in vivo*.

These findings complement ADAR1p150’s role in regulating PKR, which is activated after MDA5 sensing of unedited dsRNA ^15^. However it was not clear to what extent the type I IFN response was required for PKR activation. In the case of OAS-RNaseL, as we now describe, it is only activated after type I IFN induced signalling and transcription. This highlights that OAS proteins sense immunogenic cellular RNAs when these are not protected by ADAR1p150 binding. Our data help explain reported differences in the outcomes of RNASEL loss between human cell lines and murine models ^82, 83^. The cellular RNA sensor/binding protein repertoire - MDA5, PKR, ZBP1, OAS, RNaseL - and the nature and strength of type I IFN activation (often intrinsic in cancer cells) determine responses to ADAR1 loss. This model also explains ADAR1 and OAS-RNaseL roles in responses to demethylating agents ^81, 84^. Notably, RNaseL-deficient mice show reduced effects to radiation exposure and better recovery, suggesting ADAR1p150’s protective role may extend even when MDA5 and PKR are present ^85^. We found no role for ZBP1 in this response, despite links to ADAR1 deficient phenotypes, suggesting ZBP1’s role may be context dependent or, alternatively, fully dependent on PKR ^61, 62, 63, 64, 65, 66^.

Our analysis expands ADAR1p150’s known *in vivo* functions, positioning it as a pleiotropic suppressor of immunogenicity of cellular derived dsRNA. Editing by ADAR1p150 prevents MDA5 activation, but MDA5 can still bind unedited dsRNA even when ADAR1p150 is present ^5^. PKR and OAS engagement depends on ADAR1p150’s RNA binding, not editing, and OAS activation requires type I IFN-induced transcription (**Supplemental Figure S11B**). This study identifies that there is a distinct set of immunogenic cellular derived RNAs that are induced by type I IFN, even under sterile conditions. The levels of ADAR1p150 protein, balanced by the levels of both positive and negative regulators of other cytoplasmic dsRNA sensors ^86^, regulate the threshold level of dsRNA available at homeostasis and upon tonic type I IFN signaling to determine the cellular outcome. If the levels of cellular dsRNA are within the editing and binding capacity of ADAR1p150, the cell remains tolerant to these self-derived dsRNAs. However, once the levels of cellular dsRNA exceed those that ADAR1p150 can engage, the dsRNA become available for other dsRNA sensors such as PKR and OAS proteins to sense. ADAR1p150 activity and protein levels thus define a set point for tolerance to self dsRNAs.

## Materials and Methods

### Ethics statement

All animal experiments were approved by the St Vincent’s Hospital Melbourne Animal Ethics Committee (AEC#016/20 and #012/23) and the Hudson Institute of Medical Research Animal Ethics Committee (MMCB/2024/16).

### Mouse lines

*Adar^E861A/+^* (*Adar1^E861A/+^*; MGI allele: Adar^tm1.1Xen^; MGI:5805648), *Adar^-/-^* (*Adar1^-/-^*; MGI allele: Adar^tm2Phs^; MGI:3029862), *Adar1p150^-/-^* (p.L196C*fs*X6), *Adar1^P195A/+^*, *Ifih1^-/-^* (Ifih1^tm1.1Cln^), *Eif2ak2^-/-^* (Eif2ak2^tm1Cwe^; *Pkr^-/-^*; MGI:2182566) and *Rosa26*-CreER^T2^ (Gt(*ROSA*)26^Sortm1(cre/ERT2)Tyj^) mice have all been previously described ^5, 14, 15, 18, 19, 34^. All mice were on a backcrossed C57BL/6 background and genotyped as previously described. Where used, C57BL/6 background controls were heterozygous CD45.1/CD45.2 donor (bred internally at St Vincent’s Institute).

### Hematopoietic analysis

At the indicated ages peripheral blood (PB) samples were obtained via retro-orbital or submandibular bleeding. For analysis of bone marrow, spleen and thymus composition, PB was collected, and mice then euthanized. Femurs were flushed (2 femurs in 2mL PBS/2%FBS), spleens (5mL PBS/2%FCS) and thymus (2mL PBS/2%FCS) crushed, and single cell suspensions were prepared in PBS containing 2% FBS. Spleen and thymus were weighed and crushed through 40μM cell strainers (BD). PB, bone marrow, spleen, and thymus suspensions were counted on a Sysmex KX21 or XN-350 hematological analyzer (Sysmex Corp, Japan). Antibodies against murine B220 (APC), CD11b/Mac-1 (APC-ef780), Gr1 (PE), F4/80 (BV650), CD4 (eFluor450) and CD8a (PerCP-Cy5.5), CD45.1 (PE-Cy7), CD45.2 (AF700 or FITC), Sca-1 (PerCP-Cy5.5), c-Kit (APC-eFluor780/SuperBright600), CD150 (PE/PE-Cy7), CD48 (PE-Cy7/BV421), CD135 (PE-Cy5), CD34 (eFluor660), CD16/32 (eFluor450) and biotinylated antibodies (CD2, CD3e, CD4, CD5, CD8a, B220, Gr-1, CD11b/Mac1, Ter-119) were used. The biotinylated antibodies were detected with streptavidin-conjugated Brilliant Violet-605 or strep-FITC ^5, 87, 88^. Cells were acquired on a BD LSRII Fortessa or Cytek Aurora and analysed with FlowJo software Version 9 or 10.0 (Treestar).

### HSC and progenitor isolation

The femur, tibiae and iliac crest was isolated from male *A1+/+* and *A1p150-/-* triple mutant mice (all male; >8 weeks of age; co-housed littermates). These were cleaned of any muscle, crushed sterilely in a mortar and pestle, then filtered through 40mm cell strainers. The hematopoietic cells were then overlaid on a Ficoll-hypaque (Sigma Aldrich) layer and the mononuclear fraction enriched. Cells were washed in PBS containing 2% FCS and then stained at a concentration of 5 x 10^7^/mL with pre-titred concentrations of biotin lineage cocktail (anti-CD2, CD3, CD4, CD5, CD8, CD11b, Gr-1, Ter119, B220), Sca-1 PerCP-Cy5.5, c-Kit SB600, CD135 PE-Cy5 or CD135 PE, CD48 BV421, CD150 PE-Cy7 and CD34 eF660 or CD34 AF647. Following staining for 30 minutes in the dark on ice, the cells were washed in PBS without serum and then stained with Aqua live/dead stain (Invitrogen) and streptavidin BV786. After staining for 30 minutes in the dark on ice, the cells were washed in PBS/2% FCS and resuspended for sorting using a BD FACS Aria Fusion. One step compensation beads were used for compensation settings. The following populations were isolated: long-term hematopoietic stem cells (LT-HSC), phenotype Lin-cKit+ Sca-1+ CD135-CD150+ CD48-CD34-; MPP1, phenotype Lin-cKit+ Sca-1+ CD135-CD150+ CD48-CD34+; short-term HSCs (ST-HSC), phenotype Lin-cKit+ Sca-1+ CD135-CD150-CD48-; and MPP4, phenotype Lin-cKit+ Sca-1+ CD135+CD150-CD48+. Cells were isolated using a BD AriaIII (BD Bioscience).

### Bone marrow transplantation

Bone marrow transplants were performed into lethally irradiated (2 x 5Gy; 3hr apart; Gammacell Irradiator) female B6.SJL-Ptprca^Pep3b/BoyJArc^ recipients (from WEHI Animal Facility, Melbourne; CD45.1 expressing, 3–4 recipients/donor/transplantation cohort). For non-competitive transplants, donor bone marrow (CD45.2 expressing) was collected from femurs of 1 male donor and 1 female donor per genotype and these were transplanted separately into 3 recipients per donor; total of 6 recipients per genotype. Each recipient received 5 x 10^6^ whole bone marrow cells by intravenous injections. For competitive transplantation, 1 x 10^6^ whole bone marrow cells from a donor of each genotype was mixed with 1 x 10^6^ whole bone marrow cells from a heterozygous CD45.1/CD45.2 donor (bred internally at St Vincent’s Institute) was used as competitor. Recipient animals received antibiotics (Enrofloxacin; Baytril) in the drinking water for 3 weeks after irradiation. Animals were monitored for chimerism by FACS using peripheral blood at 5 weeks, 12 weeks and 20 weeks post-transplant and then analysis of bone marrow, spleen and thymus undertaken upon completion. Antibodies against murine B220 (APC), CD11b/Mac-1 (APC-ef780), Gr1 (PE), F4/80 (BV650), CD4 (eFluor450) and CD8a (PerCP-Cy5.5), CD45.1 (PE-Cy7) and CD45.2 (AF700 or FITC) were used. Cells were acquired on a BD LSRII Fortessa and analysed with FlowJo software Version 9 or 10.0 (Treestar).

### *In vivo* anti-IFNAR1 monoclonal antibody treatment

Mice of the indicated genotypes were identified and aged until at least 8 weeks of age. Male and female mice were used and co-housed where possible. Mice were weighed and treatment commenced with anti-IFNAR1 monoclonal antibody (MAR1-5A3; isotype mouse IgG_1_; Leinco Technologies Inc) or anti-mouse IgG_1_ control (Isotype Control; Leinco Technologies Inc) ^55, 56^. Animals were weighed before each dose and the neutralising antibody or control was administered intraperitoneally 3 times per week (Monday, Wednesday and Friday) with the first three doses being 0.5mg/dose then 0.25mg/dose for all subsequent doses. Antibodies were diluted to working concentration immediately before administration using sterile PBS.

### Mouse myeloid cell lines

Bone marrow was isolated from the femurs of adult (>8 weeks old) *Adar1^+/+^Ifih1^-/-^Eif2ak2^-/-^* (*A1+/+*; ADAR1 wild-type, MDA5 and PKR deficient; control), *Adar1^-/-^Ifih1^-/-^Eif2ak2^-/-^* (*A1-/-*; ADAR1 protein deficient, MDA5 and PKR deficient), *Adar1^E861A/E861A^Ifih1^-/-^Eif2ak2^-/-^*(*A1EA/EA*; ADAR1 editing deficient, MDA5 and PKR deficient) and *Adar1^L196C/L196C^Ifih1^-/-^Eif2ak2^-/-^*(*A1p150-/-*; ADAR1p150 protein deficient, MDA5 and PKR deficient) ^15^. After ficoll separation, cells were cultured in IMDM (Sigma) supplemented with 10% FBS (Assay Matrix; non-heat inactivated), 1% Penicillin/Streptomycin (Gibco/Thermo Fisher), 1% glutamax (Gibco/Thermo Fisher), 50ng/mL recombinant mouse stem cell factor (rmSCF, Peprotech), 10ng/mL recombinant mouse interleukin 3 (rmIL-3, Peprotech) and 10ng/mL recombinant human interleukin 6 (rhIL-6, Amgen) for 48 hrs. After 48hr in culture, 1×10^6^ cells were spin-infected at 1,100g for 90 minutes with ecotrophic packaged HOXA9 retrovirus as previously described ^15, 57, 89^ and 8ug/mL hexadimethrine bromide (Polybrene; Sigma). At 48 hrs post-infection, the cells were moved into IMDM supplemented with 10% FBS, 1% Penicillin/Streptomycin, 1% glutamax and 1% granulocyte-macrophage colony-stimulating factor (GM-CSF) conditioned medium (from BHK-HM5 cell conditioned medium). Cells were maintained in GM-CSF containing media from this point onward. Cell lines established after 3-4 weeks of culture. Proliferation and viability was monitored by trypan blue and cell counting using the Countess II (Invitrogen).

### Generation of retrovirus or lentivirus

Retrovirus was produced using transient transfection of HEK293T cells seeded in either 6-well tissue culture plates or in 10cm tissue culture dishes in 10mL complete DMEM (supplemented with 10% FCS, 1% Glutamax, 1% Pen/Strep). For 10cm plates cells were seeded at a density of 4×10^6^ cells to reach 70% confluency in 24 hours (scaled for a 6 well plate surface area). Plasmid DNA was introduced by calcium-phosphate-mediated transfection. For retrovirus, 10μg EcoPac plasmid, 10μg retroviral expression plasmid of interest, 61μL 2M Calcium Chloride and nuclease-free water up to 500μL. For lentivirus, 3.5μg pVSV-G plasmid, 6.5μg psPAX2 packaging plasmid and 10μg lentiviral expression plasmid were used. To the plasmid/CaCl_2_ solution 500μL of 2x HEPES-buffered saline (HBS) was added dropwise while vortexing. This solution was added drop by drop to the HEK293T cells, and cells incubated for 16 hours. The medium was gently replaced the next day, and cells incubated for a further 48 hours. Virus containing supernatant was collected at 48 and 72 hours in 1mL aliquots and stored at -80°C. psPAX2 was a gift from Didier Trono (Addgene plasmid # 12260; RRID:Addgene_12260) and pCAG-Eco was a gift from Arthur Nienhuis and Patrick Salmon (Addgene plasmid # 35617; RRID:Addgene_35617).

### CRISPR/Cas9 knockouts in mouse myeloid cells

To generate additional gene knockout myeloid cell lines, the cells were infected with lentivirally packaged Cas9 plus single-guideRNAs co-expressed from the lentiCRISPRv2-Puro plasmid (Addgene plasmid # 52962, or lentiCRISPRv2-Blast, a gift from Feng Zhang) by spin-infection at 1,100g for 90 minutes with 8μg/mL hexadimethrine bromide ^57^. The infected cells were then selected in IMDM supplemented with 10% FBS, 1% Penicillin/Streptomycin, 1% glutamine, 1% GM-CSF and 3μg/mL blasticidin or 0.5μg/mL puromycin (Gibco/Thermo Fisher). sgRNA sequences were taken from the Brie mouse library ^90^ and annealed oligos (IDTDna, Singapore) were cloned into the lentiCRISPRv2 vector (gift from Feng Zhang (Addgene plasmid # 52961 ; http://n2t.net/addgene:52961 ; RRID:Addgene_52961)). sgRNA sequences provided in Supplemental Dataset S2.

CRISPR gene editing was also delivered by nucleofection (Amaxa 4D, Lonza) of either recombinant CAS9 protein (ThermoScientific) complexed with crRNA/tracrRNA (IDTDna) or purified IVT Cas9 mRNA (BASE Facility, University of Queensland) co transfected with synthetic sgRNA (IDTDna). Cells were resuspended in Opti-MEM and nucleofected. Cells were cloned as described below and mutations confirmed by Sanger sequencing and western where possible.

### Generation of myeloid cell line clones

To isolate clonal myeloid cell lines, single cell suspensions were plated in methylcellulose based semi-solid media of 1% methylcellulose containing IMDM/10% FBS/1% glutamax and 1% GMCSF. Cells were plated at 500 cells were plated per 3cm tissue culture plate and incubated in a humidified chamber within a tissue culture incubator at 37°C, 5% CO_2_ and single colonies expanded over ∼7-10 days. Colonies were isolated by pipetting and placed into 96-well plates, expanded then genomic DNA isolated using Bioline genomic DNA kit (Bioline). PCR was performed using primers listed in Supplemental Dataset S2 for each gene as indicated. The PCR products were purified using the PCR cleanup kit (Bioline) and Sanger sequenced (Australian Genome Research Facility, Melbourne or MicroMon Genomics, Monash University). CRISPR-Cas9 mutations in clones were screened for frameshift mutations using the primers in Supplemental Dataset S2 and Sanger sequencing. Clones with homozygous mutations were then tested by Western blot.

### Interferon treatment of cells

Myeloid cells: Three biologically independent immortalised myeloid cell lines per genotype were plated in 12-well plates in 1mL complete media. Recombinant mouse interferon beta (IFNβ; mammalian expressed; PBL Assay Science; Cat #12405-1) was added at either 50U/mL or 500U/mL or 25U/mL recombinant mouse interferon gamma (IFNγ; Peprotech; Cat#315-05-100ug) of culture media and the cells were incubated for up to 48 hours. The cells were counted and viability assessed by trypan blue uptake using an automated cell counter (Countess, Invitrogen). Cells were additionally tested in in an Incucyte (Sartorius) as follows. Each cell line was plated at 15,000 cells/well in a 96-well plate. Cells were treated with vehicle (media) or an increasing dose of IFNβ ranging from 10U/mL to 1000U/mL. Cytotox Red dye (Sartorius) was added at time zero, the plate centrifuged at 400g for 5 minutes and then the plate incubated in the Incucyte with imaging every 4 hours for 52 hours. Results are plotted as the phase normalised Cytox red positive cells compared to time 0.

Primary hematopoietic cells: Purified LT-HSC, MPP1, ST-HSC and MPP4 were isolated from *A1+/+* and *A1p150-/-* triple mutant mice (all male; >8 weeks of age; n=4 per genotype). The isolated cells were equally divided into 4 wells containing 200-250μl of serum free expansion media (SFEM; StemCell Technologies) supplemented with 2% penicillin/streptomycin, 1% Glutamax, 50ng/mL rmSCF (Peprotech), 10ng/mL rmFlt-3L (Peprotech), 20ng/mL rhIL-6 (Amgen), 20ng/mL rhTPO (Peprotech) and 20ng/mL rmIL-3 (Peprotech). Two wells were used as control and two wells were treated with 50U/mL recombinant mouse interferon beta (IFNβ). Cells were counted on 72hrs after plating either manually using a hemocytometer and trypan blue (for LT-HSC, MPP1 and ST-HSC) or using an automated cell counter with trypan blue (for MPP4; Countess automated cell counter). For the MPP4 population only, cells were also treated with 100U/mL IFNβ for 72 hrs then counted as described.

### cDNA for overexpression

Mouse *Adar1* cDNAs (codon optimised, synthetic origin; GeneArt, Germany) were modified by subcloning of gene synthesis products (Twist Bioscience). All *Adar1* cDNA had a N-terminal in-frame eGFP fusion and a C-terminal 3xFlag tag ^68^. ADAR1p150 specific expression (designated ADAR1p150) was achieved by introduction of a consensus Kozak sequence (AGCCACC) and the p.M249A mutation to block initiation of the p110 isoform ^68^. The ADAR1p150 RNA binding mutation (designated ADAR1p150 EAA) was generated by the conversion of the KKxxK motif in each of the three dsRBD to a EAxxA motif ^67^; mutations were introduced into the parental ADAR1p150 construct by sub-cloning of gene synthesis products (IDTDna or Twist Bioscience). All cDNA were cloned into pAIP lentiviral vector (gift from Jeremy Luban (Addgene plasmid #74171; http://n2t.net/addgene:74171; RRID:Addgene_74171)) and the plasmid sequence verified (MicroMon, Monash University). The cDNA were packaged into lentivirus and used to infect the indicated myeloid cells. Cells were assessed by either microscopy (Thunder Imaging system, Leica) or flow cytometry (Cytek Aurora/BD FACS Aria Fusion). For sequencing of *Mad2l1*, RNA was extracted, cDNA generated and the sequenced by Sanger sequencing as previously described (Micromon, Monash University) ^57, 68^.

### Western blot

Western blots were performed as previously described ^57^. Primary antibodies used in this study as follows: mouse monoclonal anti-FLAG M2 (Sigma, F1804), rat monoclonal anti-mouse ADAR1 antibody (clone RD4B11; purified from in house hybridoma supernatant by MATF, Monash University)^5^, rabbit anti-MDA-5 (Cell Signaling, D74E4), mouse anti-pan-ACTIN (ThermoFisher, MS-1295-P0), rabbit anti-RNaseL (Cell Signaling, D484J; 1:1000 dilution, 80kDa expected size); anti-ZBP1 (AG-20B-0010-C100 (Zippy) supplied by Sapphire Bioscience; used at 1:1000; expected size 42-68kDa (large isoform)). Membranes were then probed with HRP-conjugated to goat anti-rat (ThermoFisher, 31470), or

### BRB-seq

Transcriptomics was undertaken on control and rmIFNβ treated myeloid cell lines (n=3 independent cell lines per genotype). Total RNA was isolated from untreated or cells treated for either 48 hours (*A1^+/+^, A1^E861A/E861A^, A1^-/-^* and *A1p150^-/-^* genotypes; n=3 independent cell lines per genotype) or 0, 0.5, 1 and 4 hours (*A1^+/+^* and *A1p150^-/-^* genotypes only; n=3 independent cell lines per genotype) with 50U/mL rmIFNβ. Cells were collected and 1mL Trisure/Trizol reagent was added and then the lysate frozen at -80C. Total RNA was extracted using a Qiagen RNeasy Mini kits (Qiagen) and then quantified using an RNA ScreenTape Assay for TapeStation Systems (Agilent) to assess the quality of the extracted RNA samples. RNA concentration was determined using a Qubit fluorometer (Invitrogen).

An equal amount of total RNA per sample (500ng total RNA per sample) was used to prepared libraries using the Mercurius BRB-seq library preparation kit (Alithea Genomics)^91^ as described by the manufacturer. This method generates multiplexed 3’ cDNA libraries. Up to 24 samples were pooled for sequencing in one library. Once the library was generated it was assessed using the D5000 DNA Screen Tape assay and Qubit fluorometer and then sent for PE150bp sequencing at Novogene (Singapore) on the Illumina platform with a target yield of 5×10^6^ reads per sample within the library.

### Bioinformatic analysis

For *A1^+/+^, A1^E861A/E861A^, A1^-/-^* and *A1p150^-/-^* genotypes; n=3 independent cell lines per genotype treated for 48hrs with 50U/mL rmIFNβ dataset.

The reads were trimmed to 28bp (_1 only). Reads were then demultiplexed into individual FASTQ files per sample using BRB-seqTools (version 1.6.1, https://github.com/DeplanckeLab/BRB-seqTools). The reads were aligned to the mouse reference genome mm10 (GRCm38) using STAR (version 2.7.10b) and counts generated (--quantMode GeneCounts). Counts were analysed in degust (https://degust.erc.monash.edu/). Genes were filtered to keep at least 5 reads, and cpm > 1 in at least 3 samples. Differential expression was calculated using edgeR-quasi-likelihood.

For 0, 0.5, 1 and 4 hours (*A1^+/+^* and *A1p150^-/-^* genotypes only; n=3 independent cell lines per genotype ) with 50U/mL rmIFNβ dataset.

Reads were demultiplexed and UMIs extracted using BRB-seqTools as described above. The nf-core/rnaseq Nextflow pipeline verion 3.10.1 ^92^ was then used to process the demultiplexed reads. The *mus musculus* ENSEMBL version 109 reference was used. Briefly, reads are trimmed using Trim Galore ^93^ and then aligned to the genome using STAR ^94^. UMI counts per gene were then found using featureCounts ^95^. Degust ^96^ was again used for differential expression calculations, using the limma-voom method ^97^. Genes with a CPM of at least 2 in at least 2 samples were used in the analysis.

Further differential expression analysis was carried out in R using the voom-limma method with a custom design matrix. This design matrix contained columns for each of the six cell lines, for time points 0.5, 1, and 4 hours (relative to 0 hours as a baseline), and a final column allowing for differential expression in the *A1p150^-/-^*genotype at 4 hours relative to the *A1^+/+^* genotype. The coefficient for this final column was used in the differential expression test.

Gene set enrichment analysis was conducted on Gene Onotology gene sets with at least 10 genes were considered. Gene sets were considered significant with FDR ≤ 0.05 by Over-Representation Analysis, found using the ClusterProfiler package.

RNAseL signature analysis: The signature of RNaseL activation by 2-5A was extracted from PMID:38823018 ^60^. The data was converted from human to 1:1 homologs with mouse genes obtained from ENSEMBL BioMart. There are 9110 genes with homologs in the RNase L differential test and present in the 48hr datasets after filtering to genes with a CPM of 2 in at least 2 samples. The signature was assessed in each cell line by computing the log2(CPM+1) values and then this was used to calculate log2 fold changes between corresponding “IFN treated” and “Ctrl” samples. The log fold changes were then compared to the log fold changes in the RNaseL signature. The comparison was performed using Spearman correlation. Statistical analysis was a one-sample t-test on the correlations for each genotype. The null hypothesis is that the correlation is zero.

## Dataset availability

GEO submission: 48hr treatment dataset

The data discussed in this publication have been deposited in NCBI’s Gene Expression Omnibus (GEO) and are accessible through GEO Series accession number GSE271978 (https://www.ncbi.nlm.nih.gov/geo/query/acc.cgi?acc=GSE271978).

The following secure token has been created to allow review of record GSE271978 while it remains in private status: **cnwlukcevzulncz**.

### Statistical Analysis

For biological experiments, the significance of results was analyzed using students t-tests, one-way or two-way ANOVA with multiple comparison corrections unless otherwise stated. All statistical tests were calculated using Prism software, with *P* <0.05 was considered significant or as indicated. Data are presented as mean ± SEM unless otherwise stated with n and P values as indicated on the figures or in the figure legends.

## Acknowledgements

The authors would like to thank E Tonkin, R Walder, M Stewart, J Howden, S Lin, P Hertzog, A Matthews, N De Weerd, N Sampaio, K Simpson for technical assistance and reagents; P Hertzog, H Kato, E Rothenburg, C Pfaller, K Quinn and J Heierhorst for comments and discussion; St. Vincent’s Hospital Bioresource’s Centre and Hudson Institute Animal Facility for care of experimental animals. St Vincent’s Institute Flow Cytometry Facility and FlowCore at Monash Health and Translational Precinct for cell sorting. The author/s acknowledge the facilities and the scientific and technical assistance of BASE at The University of Queensland for provision of IVT mRNA for Cas9. Schematic figures were made using BioRender.com.

This work was supported by: National Health and Medical Research Council Australia (NHMRC; GNT1183553 to CRW and JHF; GNT1182453 to JHF and AMC; GNT1182453 to JHF; GNT2018098 to CRW); the NHMRC Centre of Excellence in Nucleic Acid Sensing (GNT2035500; CRW); 5point Foundation (JHF); Melbourne Research Scholarship, The University of Melbourne (ZL); BASE is supported by Therapeutic Innovation Australia (TIA). TIA is supported by the Australian Government through the National Collaborative Research Infrastructure Strategy (NCRIS) program; and the Victorian State Government Operational Infrastructure Support Scheme (to St Vincent’s Institute of Medical Research and Hudson Institute of Medical Research).

## Author contributions

Conceptualization: JHF, SBH, JBL, CRW; Methodology: JHF, MFS, DI, LEP, CRW; Investigation: JHF, ST, MS, ZL, MFS, AMC, AG, LEP, CRW; Visualization: JHF, AMC, CRW; Funding acquisition: JHF, AMC, CRW; Project administration: CRW; Supervision: JHF, CRW; Writing – original draft: CRW; Writing – review & editing: JHF, ZL, MFS, DI, AMC, SBH, JBL, LEP, CRW.

## Declaration of interests

JBL is a co-founder of AIRNA Bio and a consultant for Risen Pharma. All other authors declare that they have no competing interests.

## Supplemental Information

**Supplemental Dataset S1. Transcriptomic profiling of myeloid cells treated with rmIFNβ. Supplemental Dataset S2. Primers and sgRNA sequences used in this study.**

**Dataset availability statement:**

GEO submission: 48hr treatment dataset

**Supplemental Figure 1.**
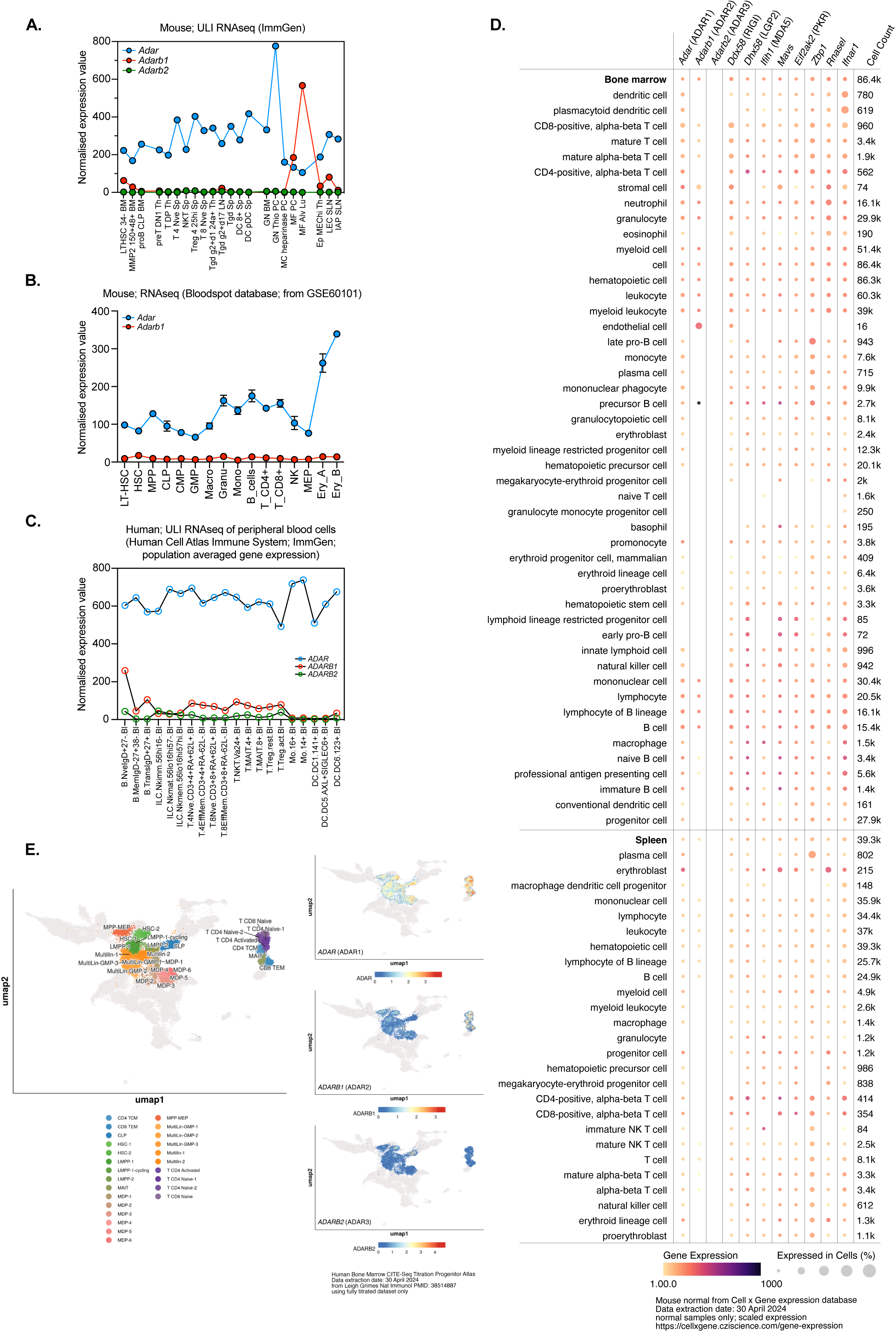
Analysis of expression of ADAR family members in mouse and human hematopoiesis. (**A**) Mouse RNA-seq datasets from ImmGen database expressed as normalised expression value. Populations: LTHSC.34-.BM - Bone Marrow 34-LTHSC/Bone Marrow 34-Long Term hematopoietic stem cells; MMP2.150+48+.BM - Bone Marrow CD150+ CD48+ MMP2; proB.CLP.BM - Bone Marrow Common Lymphoid Progenitor; preT.DN1.Th - Thymic preT DN1; T.DP.Th - Double Positive Thymocytes; T.4.Nve.Sp - Splenic Naive CD4+ T cells; T.8.Nve.Sp - Splenic Naive CD8+ T cells; Treg.4.25hi.Sp - Splenic CD25hi Tregs; NKT.Sp - Splenic Natural Killer T cells; MAIT.Sp - Mucosal-associated invariant T cells from spleen; Tgd.g2+d1.24a+.Th - Vg2+ Scart2- immature thymocytes; Tgd.g2+d17.LN - Vg2+ CD27-.LN; Tgd.Sp - Total Splenic gdT cells; NK.27-11b+.Sp - Splenic CD27-CD11b+ Natural Killer; DC.8+.Sp - Splenic CD8+; DC.pDC.Sp - Splenic Plasmacytoid; GN.BM - Bone Marrow Neutrophil; GN.Thio.PC - Thio-induced Peritoneal Neutrophil; MC.PC - Peritoneal heparinase Myeloid Cells; MF.PC - Peritoneal Macrophages; MF.Alv.Lu - Alveolar Macrophages; Ep.MEChi.Th - Thymic Medullary Epithelial cells, MHCIIhi; LEC.SLN - Subcutaneous Lymph Node Lymphatic Endothelial Cell; IAP.SLN - Subcutaneous Lymph Node (**B**). Mouse RNA-seq datasets from GSE60101 obtained from Bloodspot database; expressed as normalised expression value. Populations: LT-HSC - Long Term Hematopoietic Stem Cell (Lin-, ckit+, Sca1+, Flk2-, CD34-); HSC - Hematopoietic Stem Cell (Lin-, ckit+, Sca1+, Flk2-, CD34+); MPP - Multipotent Progenitor (Lin-, ckit+, Sca1+, Flk2+, CD34+); CLP - Common Lymphoid Progenitor (Lin-, ckit+, Flk2+, Il7R+); CMP - Common Myeloid Progenitor (Lin-, ckit+, Sca1-, CD34+, FcgRIII int); GMP - Granulocyte Monocyte Progenitor (Lin-, ckit+, Sca1-, CD34+, FcgRIII high); Macro - Bone Marrow Macrophages (B220-, CD3-, NK1.1-, F4/80+, CD115-, low SSC); Granu - Granulocytes (B220-, CD3-, NK1.1-, Gr1+, high SSC); Mono - Monocytes (B220-, CD3-, NK1.1-, CD115+, low SSC); B_cells (B220+, CD3-, CD19+); T_CD4+ -T CD4+ cells (B220-, CD3+, CD19-, CD4+, CD8); T_CD8+ - T CD8+ cells (B220-, CD3+, CD19-, CD4-, CD8+); NK - NK Cells (B220-, CD3-, CD19-, CD4-, CD8-, Terr119-, TCRbeta-, NK1.1+); MEP - Megakaryocytic erythroid progenitor (Lin-, ckit+, Sca1-, Flk2-, CD34-); Ery_A - Erythrocytes A (B220-, CD3-, Ter119+, CD71+); Ery_B - Erythrocytes B (B220-, CD3-, Ter119+, CD71-). (**C**) Human RNA-seq of peripheral blood cells from the Human Cell Atlas Immune System and ImmGen database; population averaged gene expression expressed as normalised expression value. Populations: B.NveIgD+27-.Bl - Sorted as CD19+ IgD+ CD27- PBMCs; B.MemIgD-27+38-.Bl - Sorted as CD19+ IgD- CD27+ CD38- PBMCs; B.TransIgD+27+.Bl - Sorted as CD19+IgD+CD27- PBMCs; ILC.Nkimm.56hi16-.Bl - Sorted as CD19- CD56hi PBMCs; ILC.Nkmat.56lo16hi57-.Bl - Sorted as CD19- CD56mid CD16+ CD57-PBMCs; ILC.Nkmem.56lo16hi57hi.Bl - Sorted as CD19- CD56mid CD16+ CD57+ PBMCs; T.4Nve.CD3+4+RA+62L+.Bl - Sorted as DR- TCRb+ CD4+ CD8- CD127+ CD25- CD54RAhi CD62Lhi PBMCs; T.4EffMem.CD3+4+RA-62L-.Bl - Sorted as DR- TCRb+ CD4+ CD8- CD127+ CD54RA- CD62L- PBMCs; T.8Nve.CD3+8+RA+62L+.Bl - Sorted as DR- TCRb+ CD4- CD8+ CD127+ CD25- CD54RAhi CD62Lhi PBMCs; T.8EffMem.CD3+8+RA-62L-.Bl - Sorted as DR- TCRb+ CD4- CD8+ CD127+ CD54RA- CD62L- PBMCs; T.NKT.Va24+.Bl - Sorted as CD19- CD3+ XXX PBMCs; T.MAIT.4+.Bl - Sorted as CD19- CD3+ CD4+ CD8- Va72.2+ CD161dull PBMCs; T.MAIT.8+.Bl - Sorted as CD19- CD3+ CD4- CD8+ Va72.2+ CD161dull PBMCs; T.Treg.rest.Bl - Sorted as CD19- CD4- CD25+ CD127- CD45RA+ PBMCs; T.Treg.act.Bl - Sorted as CD19- CD4- CD25+ CD127- CD45RA- PBMCs; Mo.16+.Bl - Sorted as HLA.DR+ CD16hi CD14lo PBMCs; Mo.14+.Bl - Sorted as HLA.DR+ CD16lo CD14hi PBMCs; DC.DC1.141+.Bl - Sorted as HLA.DR+ CD141+ CD16- CD14- PBMCs; DC.DC5.AXL+SIGLEC6+.Bl- Sorted as HLA.DR+ CD141- CD16- CD14- AXL+ SIGLEC6+ PBMCs; DC.DC6.123+.Bl - Sorted as HLA.DR+ CD141- CD16- CD14- AXL- SIGLEC6- CD123+ CD11c- PBMCs. (**D**) Data from CellxGene single cell analysis of murine cells of the indicated population and gene. (**E**) *ADAR* (ADAR1), *ADARB1* (ADAR2) and *ADARB2* (ADAR3) expression in human bone marrow derived CITE-seq atlas from PMID: 38514887. Expression in the HSPC and T cell populations is shown.

**Supplemental Figure 2:**
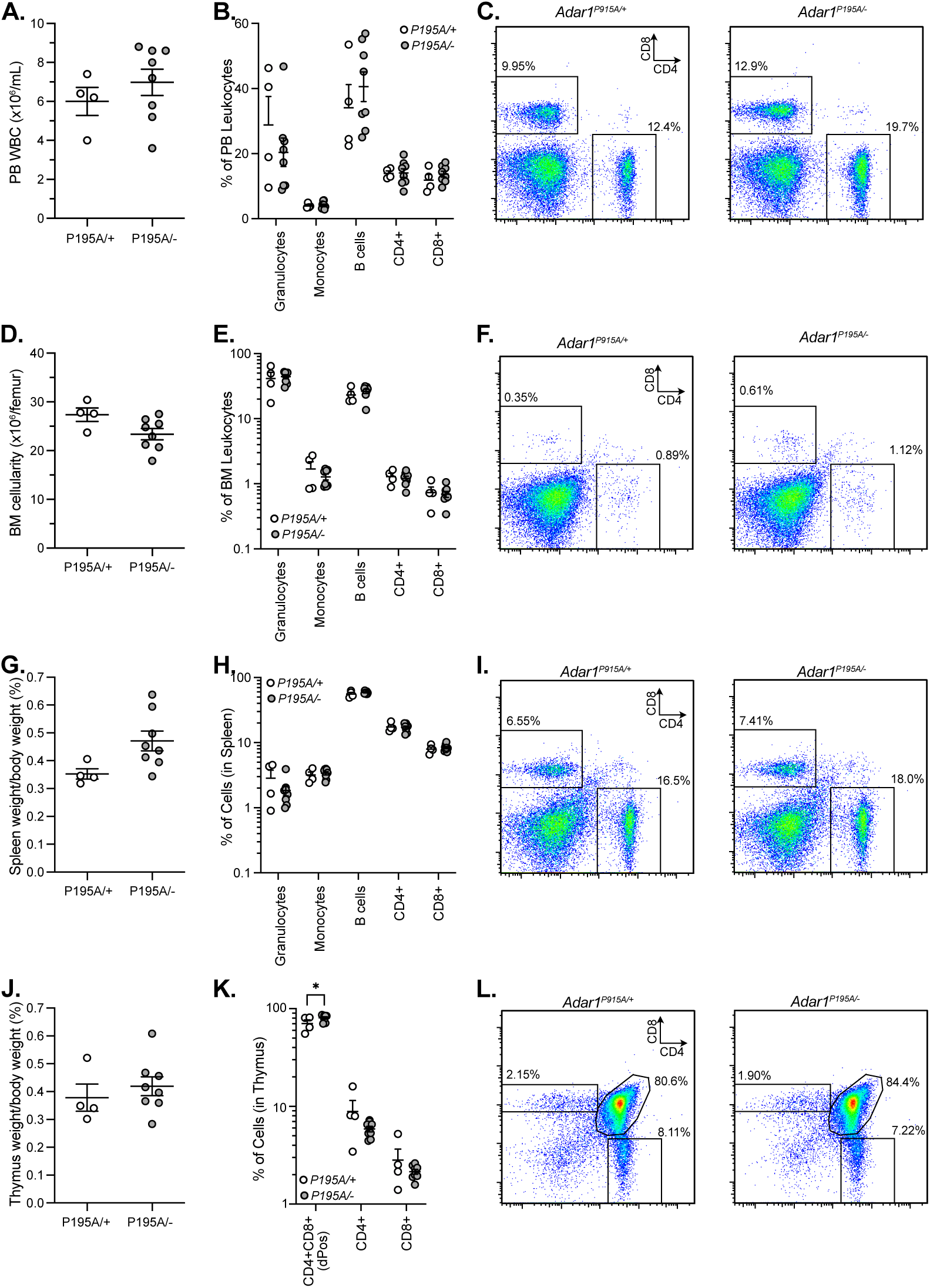
Mutations in the ADAR1p150 specific Zα domain do not impact T cell development or peripheral survival. Analysis of *Adar1^P195A/+^* (n=4; 2 male/2 female) and *Adar1^P195A/-^*(n=8; 4 male/4 female) cohorts; *ADAR* p.P195A is the most common variant in human ADAR1 reported in patients with Aicardi-Goutières’s Syndrome. Peripheral blood (**A**) leukocyte number; (**B**) lineage distribution and (**C**) representative FACS plots of T cells in the PB. Bone marrow (**D**) leukocyte number per femur; (**E**) lineage distribution of the leukocytes and (**F**) representative FACS plots of BM T cells. (**G**) Spleen weight as a proportion of body weight; (**H**) lineage distribution of splenic leukocytes and (**I**) representative FACS plots of splenic T cells. (**J**) Thymus weight as a proportion of body weight; (**J**) The proportion of cells of the indicated cell surface phenotype in the thymus and (**L**) representative FACS plots of CD4/CD8 staining of the thymus. Each data point represents an individual animal, mean ± SEM as indicated. Statistics calculated in Prism; *p<0.05.

**Supplemental Figure 3:**
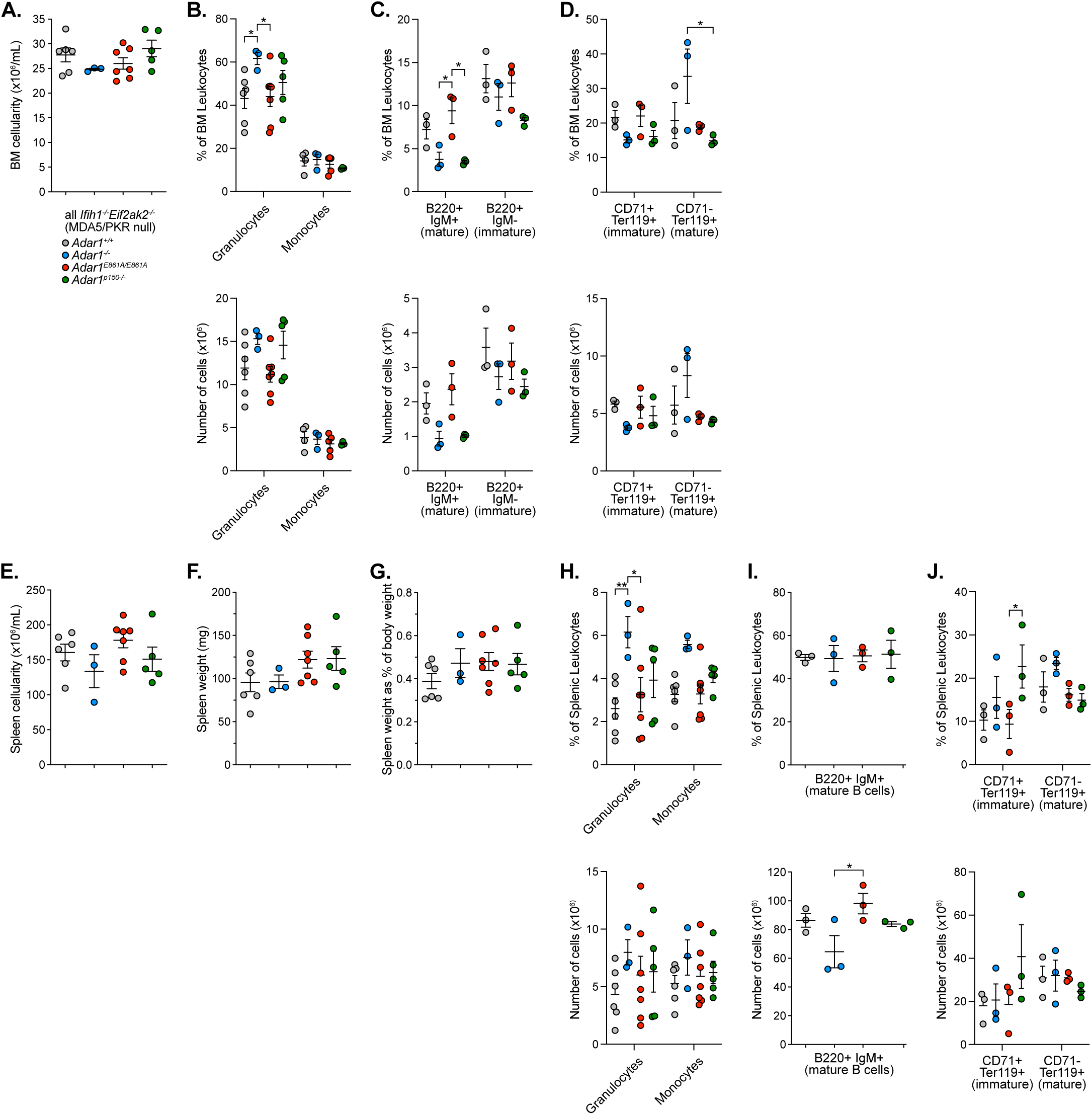
Hematopoietic cell populations in the bone marrow and spleen of the indicated genotypes. (**A**) Bone marrow cellularity per femur and percentage (upper panels) and number per femur (lower panels) of (**B**) Granulocytes (CD11b+/Gr-1+) and macrophages (CD11b+/F4/80+), (**C**) B lymphocytes (mature = B220+/IgM+; immature = B220+IgM-) cells and (**D**) erythroid (immature = CD71+/Ter119+; mature = CD71-/Ter119+) cells. Spleen (**E**) cellularity and (**F**) weight; and (**G**) spleen weight as a proportion of body weight. Percentage (upper panels) and number per spleen (lower panels) of (**H**) Granulocytes (CD11b+/Gr-1+) and macrophages (CD11b+/F4/80+), (**I**) B lymphocytes (mature = B220+/IgM+) cells and (**J**) erythroid (immature = CD71+/Ter119+; mature = CD71-/Ter119+) cells. All data from male mice 10-12 weeks of age; n per genotype as indicated in each plot with each data point and individual animal, mean ± SEM as indicated; Statistics: 1-way or 2-way ANOVA with Tukey’s multiple comparison test correction; *P<0.05; **P<0.01. All statistical tests calculated in Prism.

**Supplemental Figure 4:**
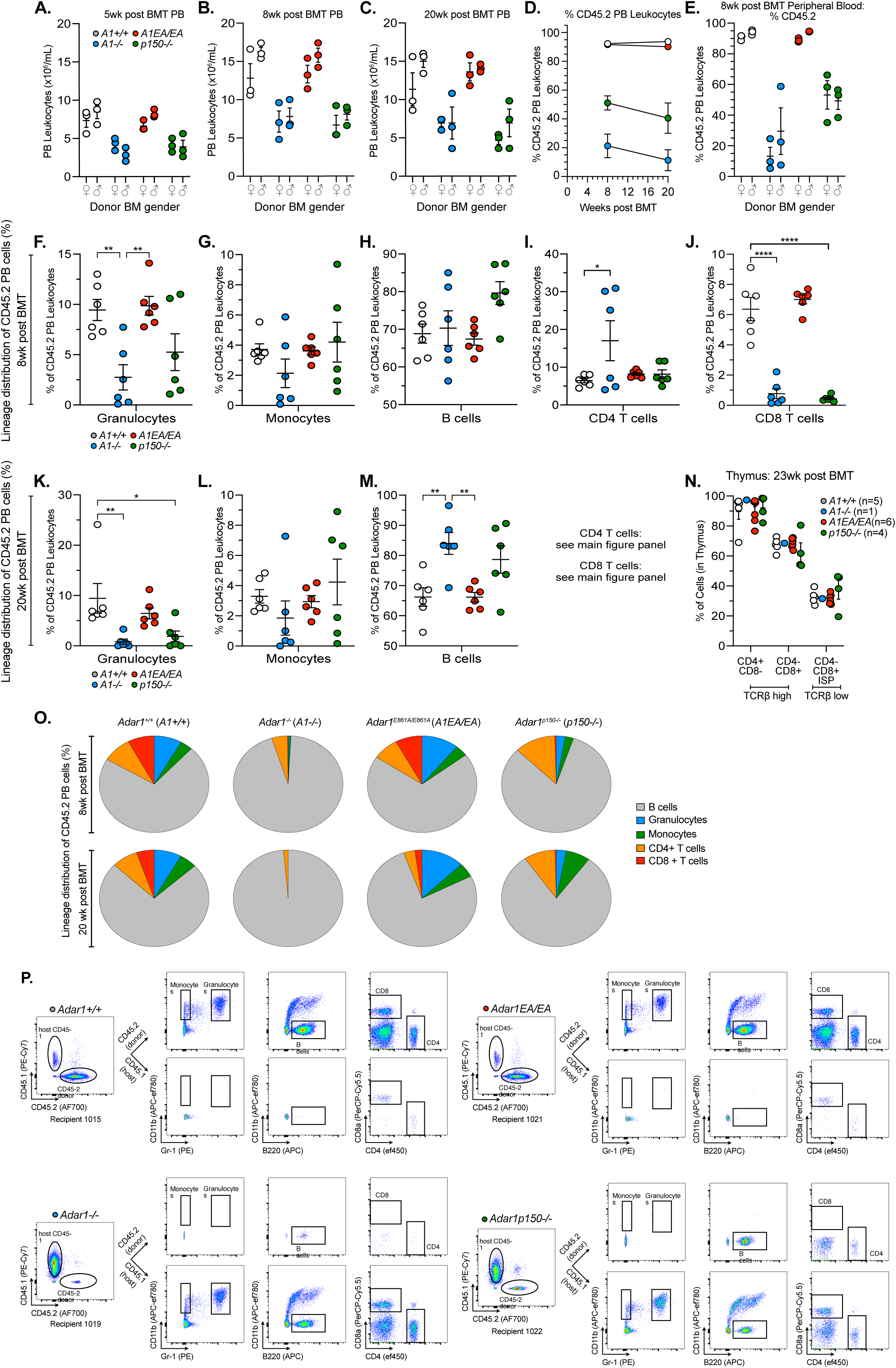
Non-competitive bone marrow transplant data. Peripheral blood leukocyte number by genotype and sex of donor at (**A**) 5 weeks, (**B**) 8 weeks and (**C**) 20 weeks post bone marrow transplant; each dot indicates an individual recipient. (**D**) Percentage of CD45.2 (donor derived) leukocytes in the peripheral blood and (**E**) CD45.2 chimerism of the PB at 8 weeks post-transplant by donor sex. (**F-J**) Lineage distribution of the CD45.2% donor cells in the peripheral blood at 8 weeks post-transplant. (**K-M**) Lineage distribution of the CD45.2% donor cells in the peripheral blood at 20 weeks post-transplant. (**N**) Percent of cells of the given TCRβ phenotype in the thymus at 23 weeks post-transplant analysis. (**O**) Pie charts of the average lineage distribution of the donor derived peripheral blood cells across each genotype. (**P**) Representative FACS plots of peripheral blood leukocytes at 20 weeks post-transplant from an *Adar1^+/+^*, *Adar1^-/-^*; *Adar1^E861A/E861A^* and *Adar1^p150-/-^*. Statistics: 1-way or 2-way ANOVA with Tukey’s multiple comparison test correction; *P<0.05; **P<0.01. All statistical tests calculated in Prism, mean ± SEM as indicated.

**Supplemental Figure 5:**
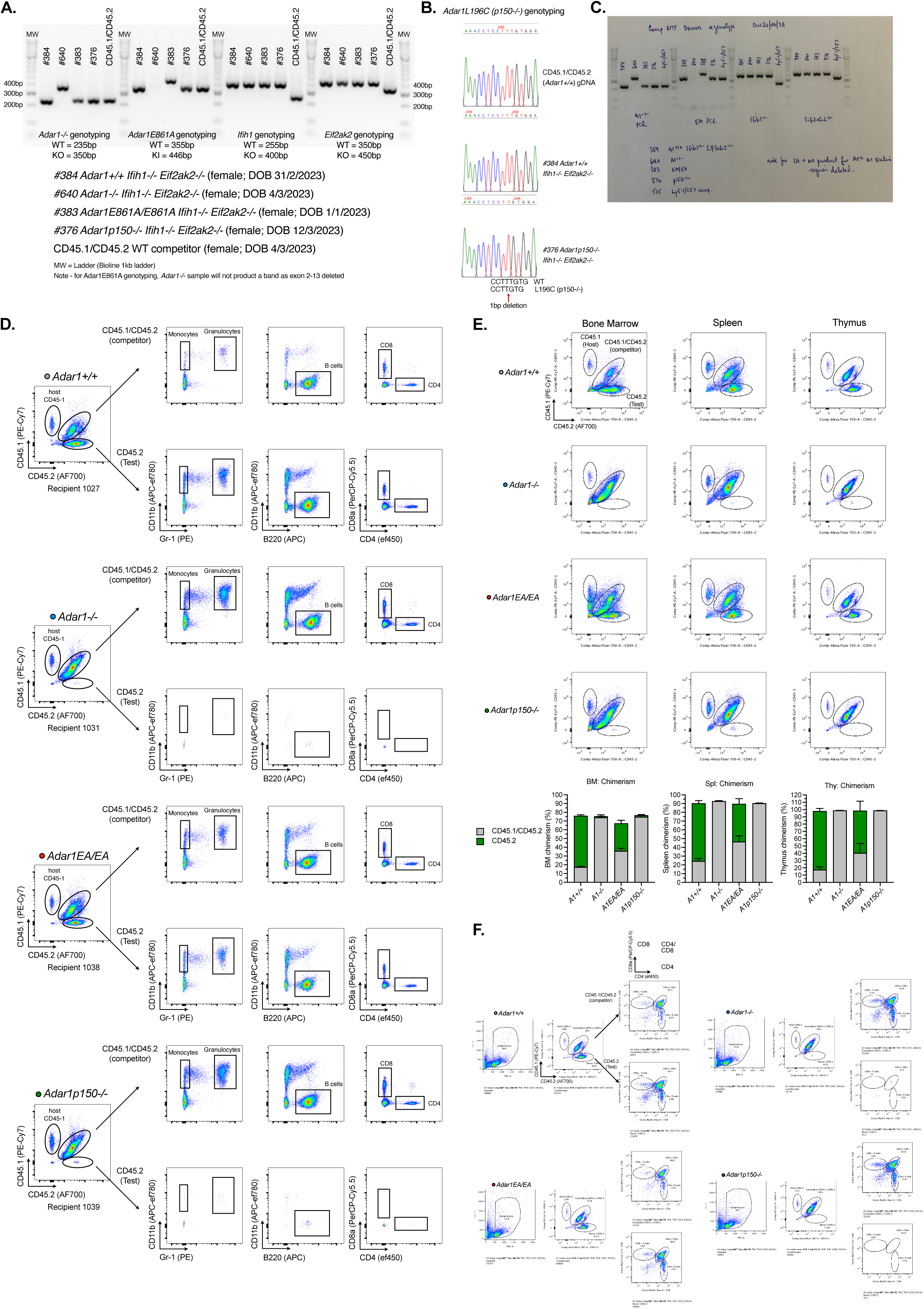
Competitive bone marrow transplant data. (**A**) Genotyping of the animals used as bone marrow donors for competitive transplants. (**B**) Sanger sequencing traces to genotype the *Adar1^p150-/-^* allele (*Adar1^L196C/L196C^*; *A1^p150-/-^*). (**C**) Uncropped image of panel A. (**D**) Representative FACS plots of peripheral blood from recipients of the indicated donor BM genotype. (**E**) Representative FACS plots of bone marrow, spleen and thymus donor and competitor cell chimerism measured at end point. (**F**) Representative FACS plots of thymocytes showing CD4 and CD8 cell population from the donor and competitor cells.

**Supplemental Figure 6:**
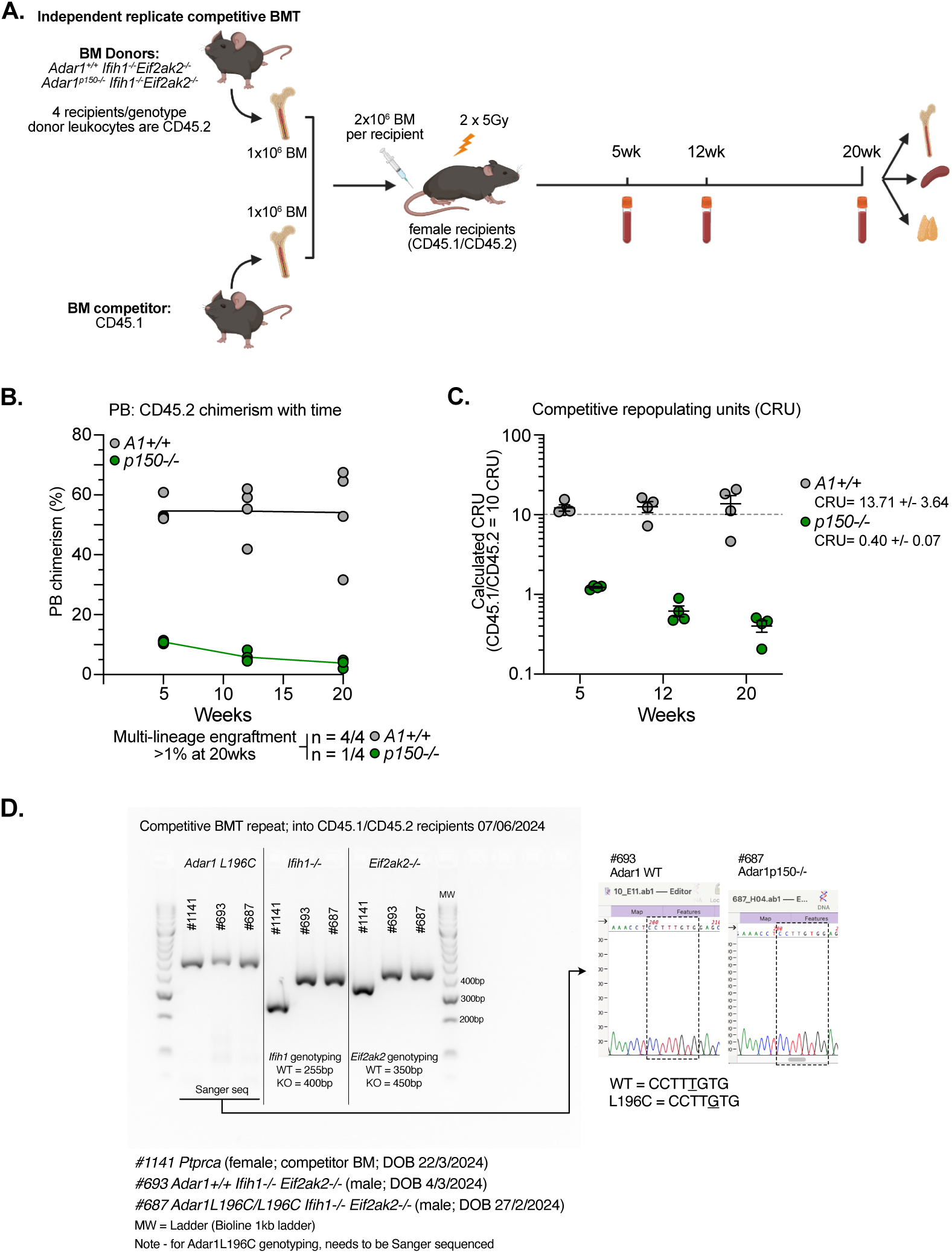
Independent replication of competitive bone marrow transplant for *A1^+/+^* and *A1^p150-/-^* bone marrow cells. (**A**) Schematic outline of the competitive transplant experiment. (**B**) Percentage CD45.2 chimerism of peripheral blood leukocytes over time for each genotype; each dot represents an individual recipient. (**C**) Calculated competitive repopulating units (CRU) for each genotype at each time point indicated. (**D**) Genotyping of the animals used as bone marrow donors for competitive transplants and Sanger sequencing traces to genotype the *Adar1^p150-/-^* allele (*Adar1^L196C/L196C^*; *A1^p150-/-^*);, mean ± SEM as indicated.

**Supplemental Figure S7:**
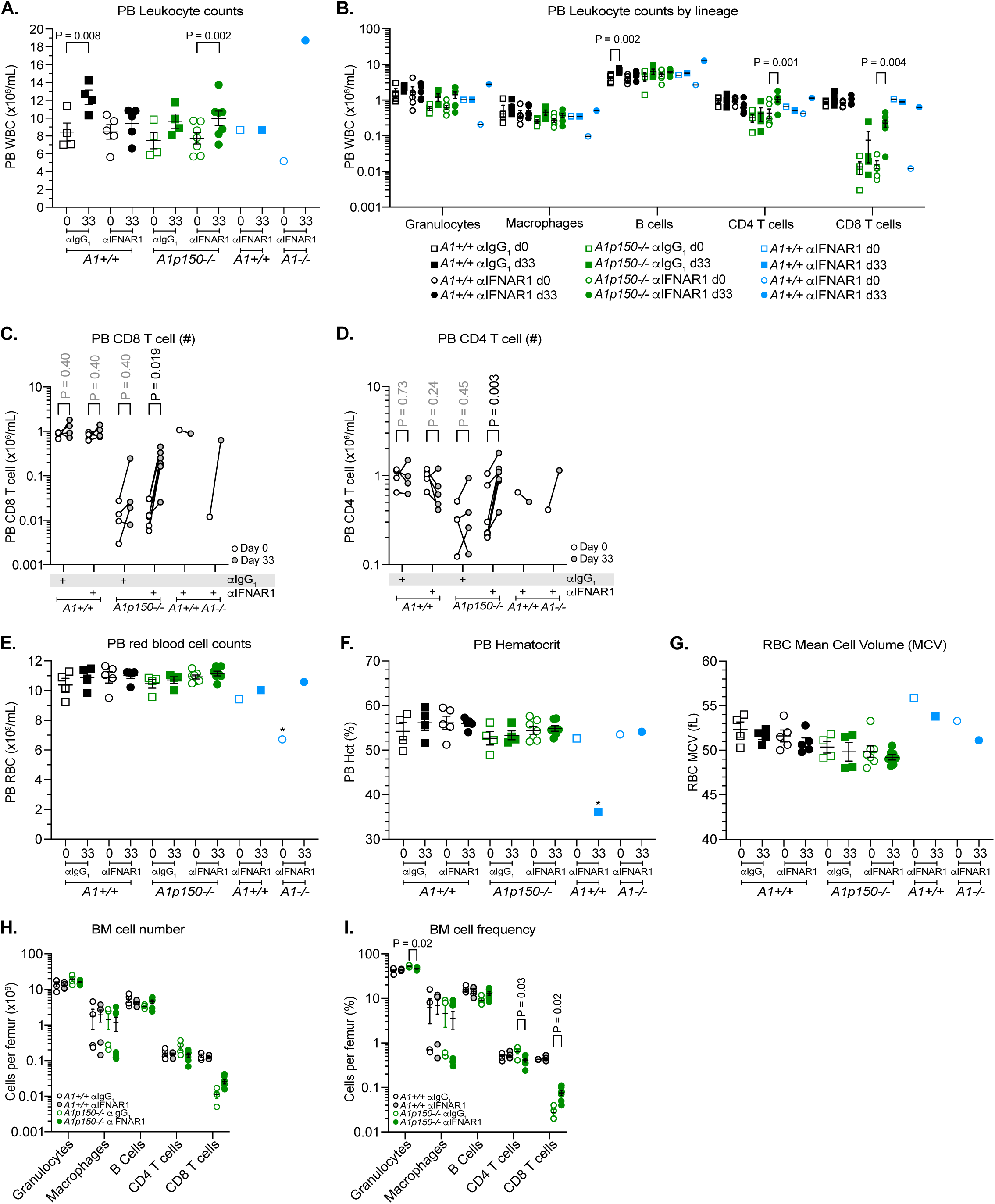
Extended data from *in vivo* treatment with αIFNAR1 monoclonal antibody. (**A**) Peripheral blood leukocyte counts at day 0 and day 33 of treatment with the indicated monoclonal antibody. (**B**) Lineage distribution of the peripheral blood leukocytes at day 0 and day 33 of treatment with the indicated monoclonal antibody. (**C**) Number of peripheral blood CD8 and (**D**) CD4 T cells in each genotype and treatment cohort; Data from each individual animal is linked by a line and all animals across both replicates pooled. Peripheral blood (E) red blood cells, (F) hematocrit and (G) mean red cell volume at day 0 and day 33 of treatment with the indicated monoclonal antibody. (H) Number of cells of each lineage per femur; (I) Frequency of cells of each lineage per femur. Each data point represents as individual animal and mean ± SEM as indicated and all animals across both replicates pooled, Statistical significance determined paired two-tailed t-tests (panel A and B; comparing time d0 to d33 for each genotype/treatment) or by multiple paired T test with FDR correction for multiple comparisons.

**Supplemental Figure S8:**
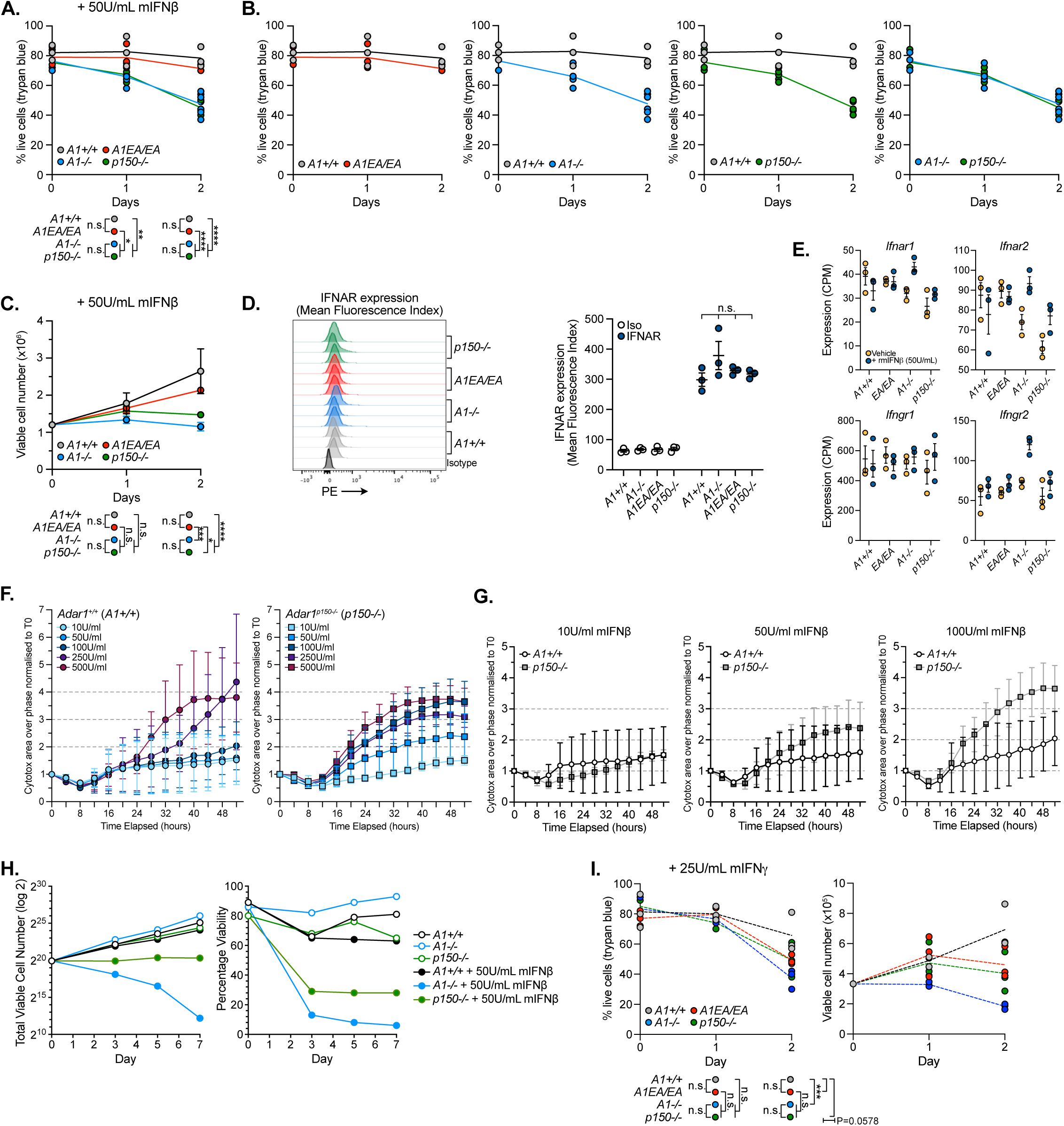
Myeloid cell lines lacking ADAR1p150 protein show hypersensitivity to low dose type I IFN. (**A**) Replicate independent experiment to data in Figure 6 on A1 triple myeloid cells treated with 50U/mL rmIFNβ; data presented as percentage of live cells (compared to untreated time 0; trypan blue staining) following treatment for 48 hours; each individual cell line indicated by a dot; statistics calculated using 2-way ANOVA with Tukey’s multiple comparison correction. (**B**) Comparisons by indicated genotype (using same data as panel A). (**C**) Viable cell number of each indicated genotype treated with 50U/mL rmIFNβ for 48 hours; each individual cell line indicated by a dot; statistics calculated using 2-way ANOVA with Tukey’s multiple comparison correction. (**D**) Cell surface IFNAR1 expression on the myeloid cell lines compared to anti-IgG_1_κ PE isotype control; each profile is from an independent cell line of the indicated genotype and quantitation and analysis of the mean fluorescence index of IFNAR1 expression (based on analysis in FlowJo). (**E**) Transcript counts of *Ifnar1*, *Ifnar2*, *Ifngr1* and *Ifngr2* from RNA-seq of untreated and cells treated with 50U/mL rmIFNβ for 48 hours; each individual cell line indicated by a dot. Data expressed as mean ± sem normalized counts per million reads (CPM). (**F-G**) The effect of increasing doses of rmIFNβ on *A1^+/+^* or *A1^p150-/-^* cells as measured using continuous monitoring with Incucyte across time. Data show increase in cell death with dose and time as measured by Cytotox area over phase normalised area at Time 0. (**H**) Effect of 50U/mL rmIFNβ treatment on the indicated genotypes for up to 7 days; Cell counts and viability from Countess/Trypan blue. Cell lines - *A1^+/+^* = 213 (derived from mouse #213); *A1^-/-^* = 616; *A1^p150-/-^*= 207. (**I**) Treatment of cells with 25U/mL rmIFNγ. Each individual cell line indicated by a dot; statistics calculated using 2-way ANOVA with Tukey’s multiple comparison correction. ***p<0.001.

**Figure S9:**
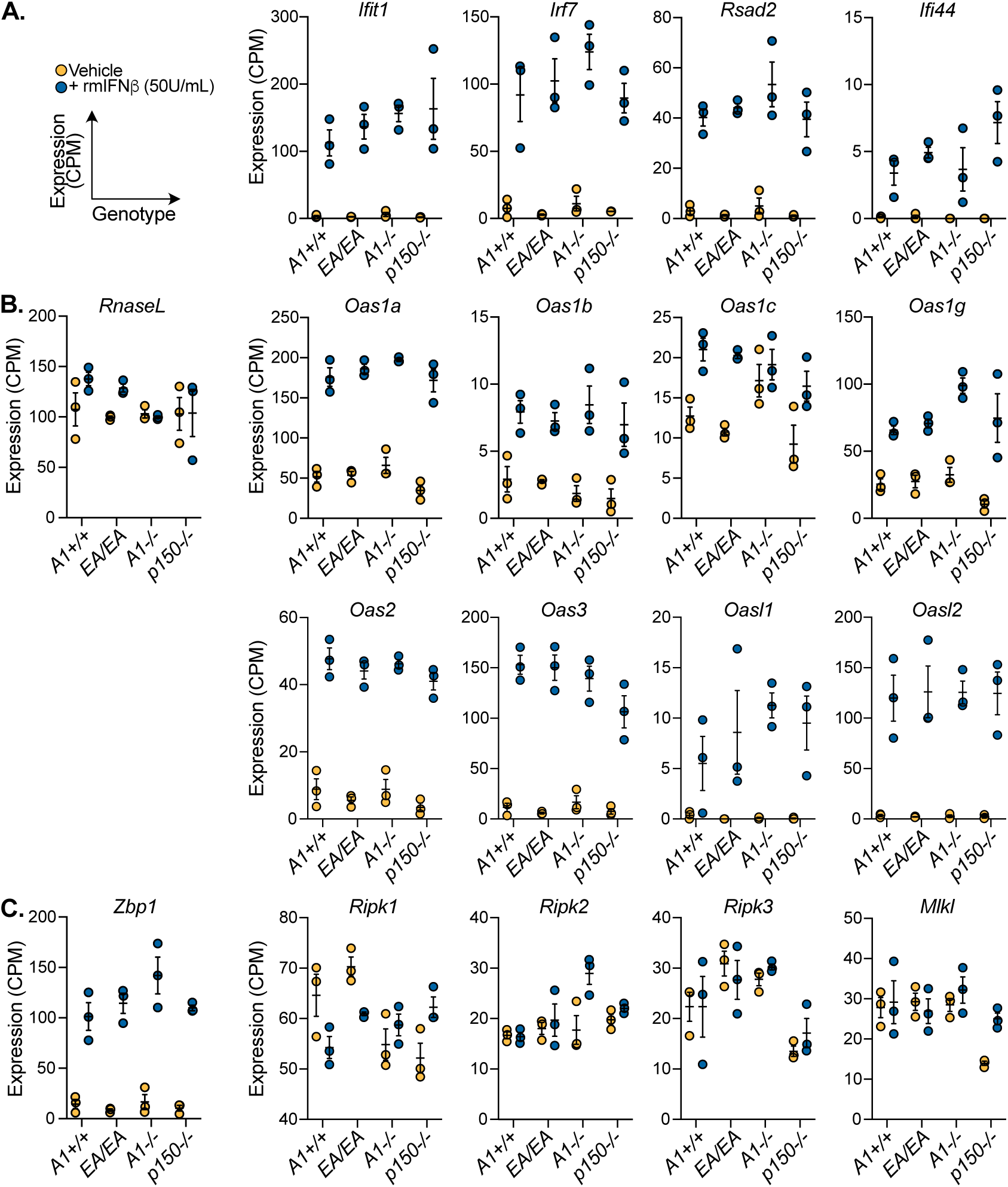
Expression of interferon stimulated genes, *RnaseL*, *Oas* genes and *Zbp1* and effectors in the myeloid cell lines. Transcript expression as measured by RNA-seq (BRB-seq) and expressed as normalized counts per million (CPM) from control or cells treated with 50U/mL rmIFNβ for 48 hours. (**A**) Expression of canonical interferon stimulated genes (ISGs); (**B**) *RnaseL* and the *Oas* transcripts and (C) *Zbp1* and related effectors in cells of the indicated genotypes and treatments. Each individual cell line indicated by a dot.

**Figure S10:**
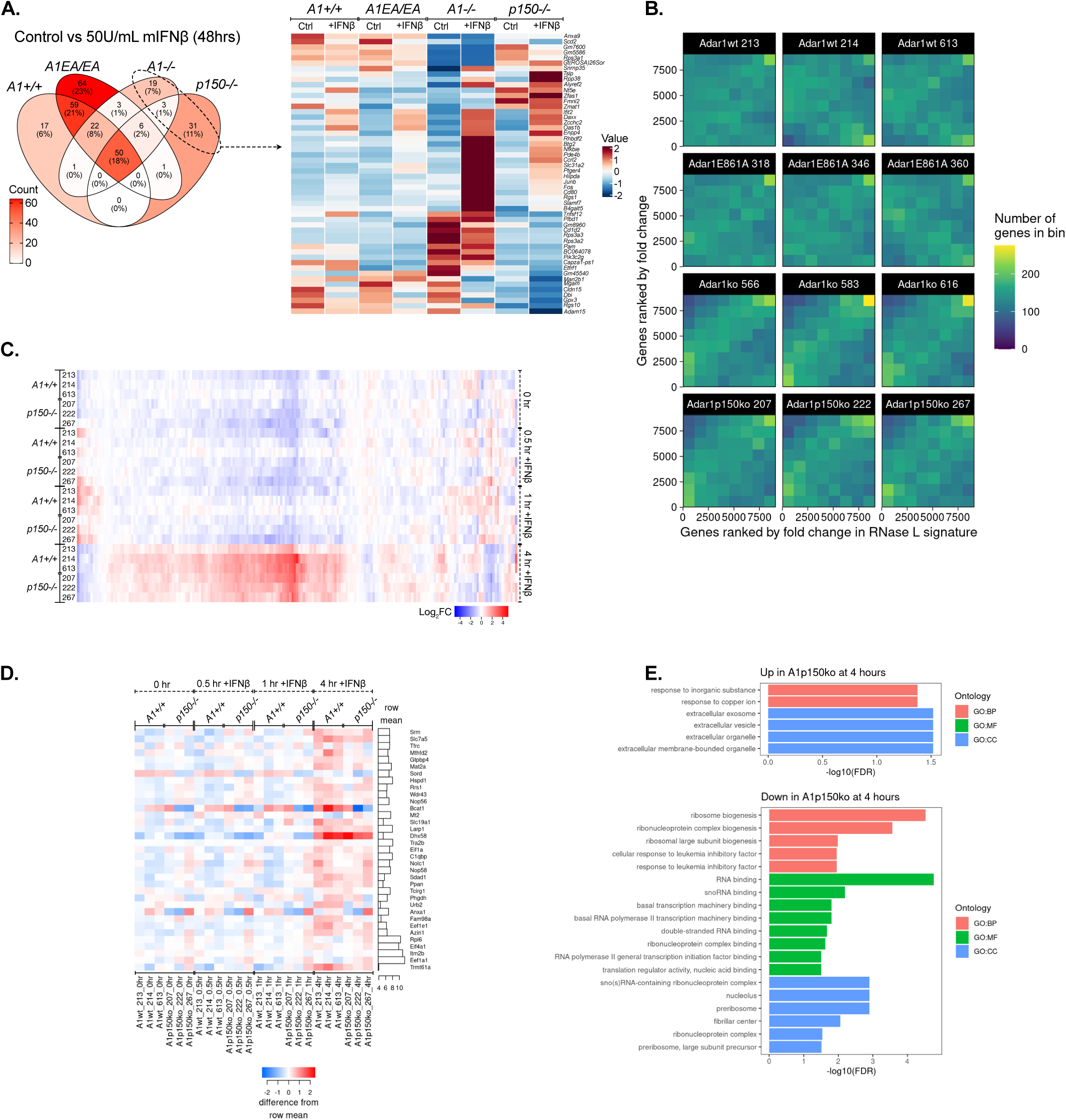
Transcriptome analysis following treatment of myeloid cells with rmIFNβ. (**A**) Overlap of the differential gene expression from RNA-seq of 48hr treated and untreated cells and a heatmap of the differentially expressed transcripts in each genotype. All genotypes n=3 independent lines per genotype; mean ± SEM as indicated. (**B**) Analysis of the 2’-5’A signature of direct RNaseL activation using Spearman correlation. (**C**) RNA-seq of time-course of *A1^+/+^*and *A1^p150-/-^* with 50U/mL IFNβ for the indicated times (0, 0.5hr, 1hr and 4hrs) normalised with the limma-voom method in Degust showing all genes LFC>1; FDR 0.05 compared to WT time 0hr. (**D**) RNA-seq of time-course of treatment of *A1^+/+^* and *A1^p150-/-^* with 50U/mL IFNβ from panel C showing differential expressed genes at 4 hours treatment shown from this analysis (no differences at 0.5 and 1hr treatment). (**E**) Pathway analysis of the time course dataset using Gene Onotology gene sets with at least 10 genes were considered. Gene sets are shown with FDR ≤ 0.05 by Over-Representation Analysis, found using the ClusterProfiler package. Numbers are log_2_ fold changes. BP: biological process, MF: molecular function, CC: cellular component.

**Figure S11:**
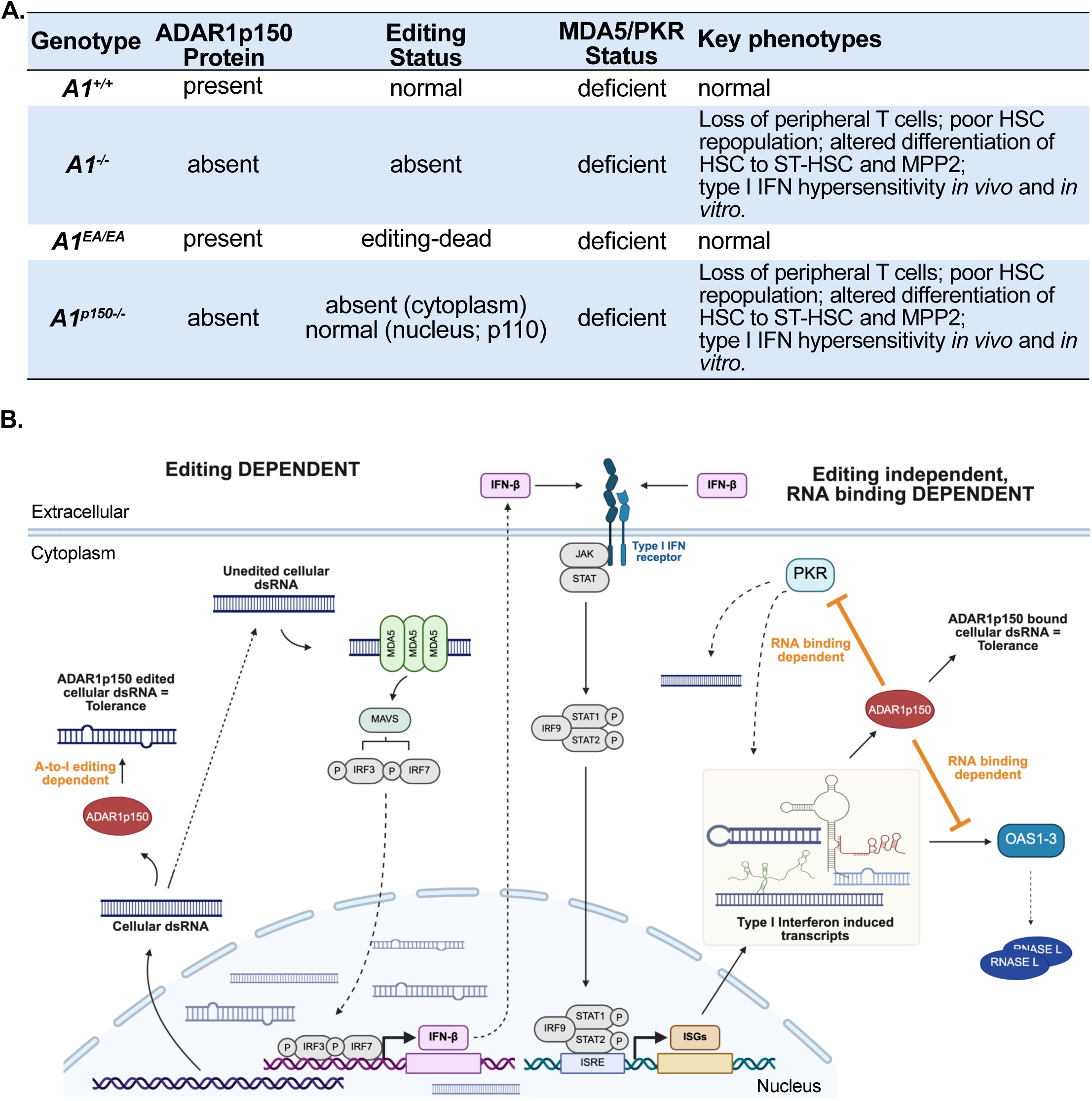
Summary of phenotypes and model. (**A**) Summary of the murine phenotypes associated with each type of mutation in ADAR1. (B) Summary of the proposed model. A-to-I RNA editing by ADAR1p150 is required to prevent MDA5 sensing of un/under-edited cellular dsRNA. MDA5 sensing initiates a MAVS dependent type I interferon response that includes production of IFNβ itself and other cellular transcripts. RNA biding, not editing, by ADAR1p150 is required to bind the cellular transcripts to prevent activation of both PKR and OAS/RNaseL. The transcripts that activate OAS/RNaseL are themselves induced by Type I Interferon.

**Figure S12:**
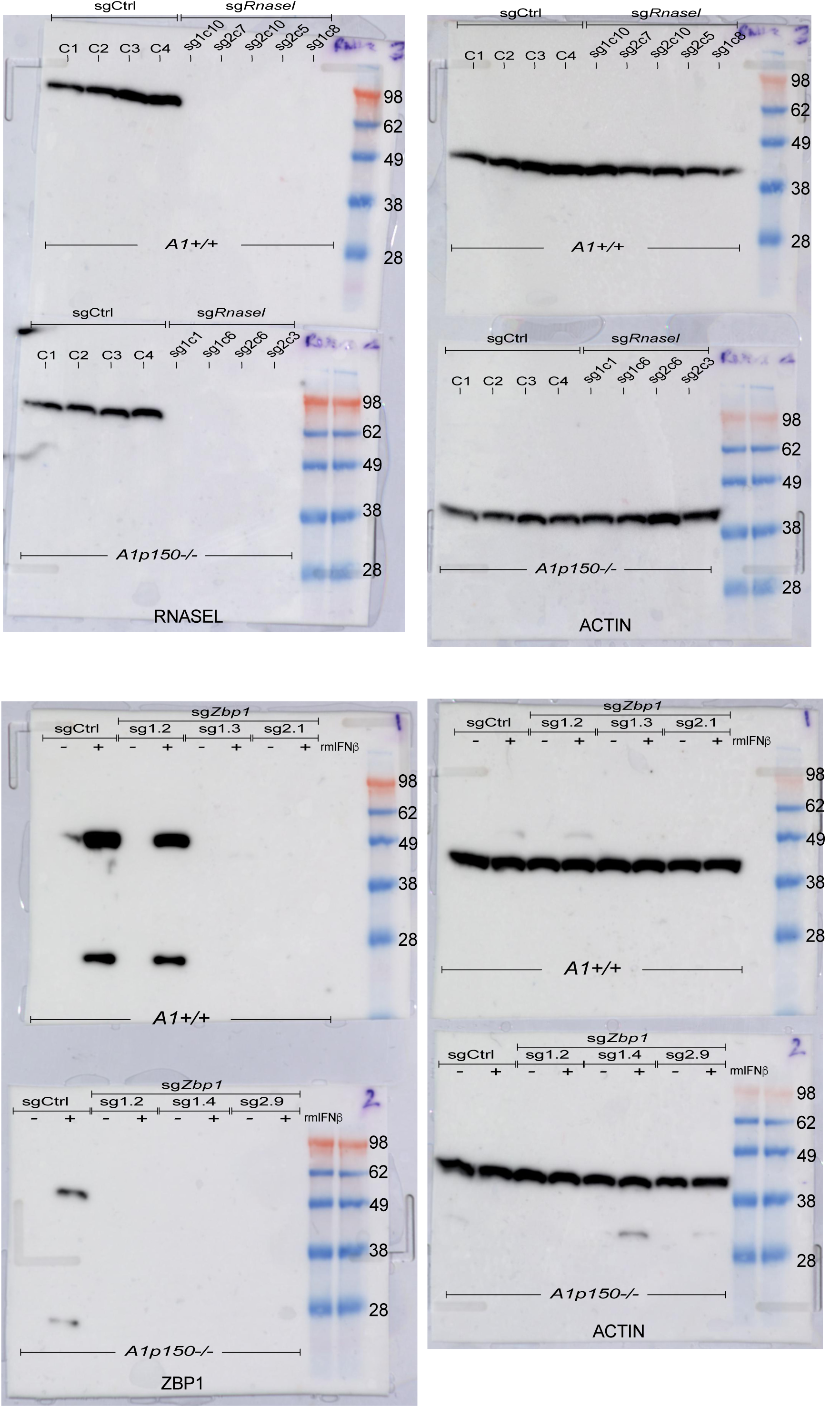
Uncropped western blots (related to. **Figure 6G and 6H)** Uncropped western blots from Figure 6G and 6H.

